# Environmental correlates of understory plant diversity and distribution in a temperate climate-change refugium

**DOI:** 10.64898/2026.10.02.756313

**Authors:** Bailey H. McNichol, Cheryl Dunn, Drew Granville, Lillian Hoffart, Brittni McGuire, Abigail Ridder, Ran Wang, Sabrina E. Russo

**Affiliations:** Wisconsin Environmental Mesonet, Wisconsin Energy Institute, University of Wisconsin-Madison, Madison, Wisconsin, USA; Department of Forestry, Michigan State University, East Lansing, Michigan, USA; School of Biological Sciences, University of Nebraska–Lincoln, Lincoln, Nebraska, USA; Department of Agronomy and Horticulture, University of Nebraska–Lincoln, Lincoln, Nebraska, USA; Bellevue University, Bellevue, Nebraska, USA; School of Natural Resources, University of Nebraska–Lincoln, Lincoln, Nebraska, USA; Department of Forest and Rangeland Stewardship, Colorado State University, Fort Collins, Colorado, USA; Center for Plant Science Innovation, University of Nebraska–Lincoln, Lincoln, Nebraska, USA

**Keywords:** climate-change refugia, diversity, ecotone, herbaceous plant, microclimate, microrefugia, topographic gradient, woody plant

## Abstract

**Aim:** Regional climate-change refugia – areas where abiotic conditions related to hydrology and topography sustain unique plant communities despite an otherwise unsuitable regional macroclimate – have been essential for maintaining biodiversity through past climatic transitions. At fine spatial scales, regional refugia can also contain microrefugia, where canopy structure and topography strongly buffer understory microclimate. We tested whether such microrefugia structure understory plant communities within a regional climate-change refugium, where sheltered, groundwater-fed canyons have sustained a biogeographically distinctive flora since the Last Glacial Maximum.

**Location:** Temperate forest in the North American Great Plains, Nebraska, USA.

**Methods:** In a 20.2-ha forest inventory plot spanning a sharp topographic gradient, we quantified diversity, composition, and distributions and of understory woody and herbaceous plants across seven functional groups along fine-scale topographic, light, and soil moisture gradients.

**Results:** Woody species density and richness were significantly higher within sheltered microrefugia in canyon bottoms and on forested slopes and strongly decreased with increasing canopy openness, whereas C_4_ grass richness was highest along the exposed, high-light forest-prairie ecotone, reflecting divergent functional group responses to microclimate buffering. Soil moisture structured richness across functional groups: woody plant and C_3_ grass richness peaked at moisture extremes, sedge richness at intermediate moisture variability, and C_4_ grass richness declined with increasing moisture variability. Both woody and herbaceous understory plants showed strong variation in composition with topography; in particular, woody seedlings and graminoids exhibited strong species turnover with increasing elevation.

**Main Conclusions:** These results demonstrate that topographically driven microrefugia support disproportionate understory biodiversity and suggest that protecting both regional refugia and finer-scale microrefugia will be critical for conserving plant communities, including species documented exclusively in microrefugia, as macroclimate continues to change.

## 1. Introduction

Ongoing climate change – characterized in temperate regions by warming temperatures, altered precipitation regimes, and increasing climate variability – is reshaping the distribution of suitable habitats for plants, creating mismatches between populations and local conditions (Davis and Shaw 2001; Hampe and Jump 2011; IPCC 2022; McNichol and Russo 2023). While many tree species are expected to track shifting climatic envelopes by migrating poleward or to higher elevations (Skov and Svenning 2004; Feeley et al. 2011; Vellend et al. 2021), understory plants (woody shrubs and herbaceous plants) face particular challenges, including more limited dispersal capacity relative to tree species (Cain et al. 1998; Van der Veken et al. 2007; Bellemare and Moeller 2014). Moreover, their smaller stature means that they may be more sensitive to finer-scale gradients in abiotic environmental conditions than larger-statured individuals (Gilliam 2007; Deng et al. 2025) that are often the focus of research in forests (Gilliam 2014; Spicer et al. 2022). Persistence in regional climate-change refugia – areas where local abiotic conditions, often driven by variation in hydrogeology and topography sustain forest communities that would otherwise be excluded by the regional macroclimate (Dobrowski 2011; Morelli et al. 2016; Stralberg et al. 2020b) – therefore represents a critical alternative to range shifting for members of the understory flora. Within regional refugia, certain areas may be more strongly buffered than others, driven by fine spatial scale variation in canopy structure, topography, and hydrology. We refer to these areas as microrefugia – areas within regional refugia where microclimate buffering is exceptionally strong in terms of decoupling understory conditions from the regional macroclimate (Dobrowski 2011; Hylander et al. 2015). Such buffering arises through the interacting effects of forest canopy shading and evapotranspiration, topographic position, and groundwater inputs, which together reduce temperature extremes and sustain soil moisture (De Frenne et al. 2019; Rita et al. 2021; Greiser et al. 2024; Koelemeijer et al. 2024). Both regional climate-change refugia and the microrefugia nested within them have been critical in maintaining regional diversity through past episodes of climatic instability (Kaul et al. 1988; Tzedakis et al. 2002; Birks and Willis 2008; Ackerly et al. 2020) and are increasingly recognized as essential biodiversity reservoirs under contemporary climate change (Morelli et al. 2020; García et al. 2022).

Plant functional groups – assemblages of species sharing traits related to growth form, physiology, and resource acquisition (Díaz and Cabido 1997; Wang et al. 2009; Douma et al. 2012; Table 1) – differ in mature size, leaf morphology, photosynthetic pathway, and stress tolerance (Díaz 2001; Wullschleger et al. 2014). Therefore, some groups are likely to sort into microrefugia where buffering is stronger while others persist in more exposed habitats where buffering is weaker (Murphy et al. 2016). Plant species richness varies strongly along broad environmental gradients (Kreft and Jetz 2007), yet how the fine-scale gradients defining microrefugia structure diversity among functional groups is poorly resolved, limiting predictions of which components of understory diversity are most at risk from macroclimatic change (Gehlhausen et al. 2000; Harrison et al. 2006; Moeslund et al. 2013). While woody plant density and diversity respond strongly to topography and light availability (Kobe 1999; Magee et al. 2021; Abiem et al. 2023), how herbaceous functional groups respond to these gradients remains poorly characterized, particularly in studies where microclimate is directly measured alongside plant diversity (Xu et al. 2000; Small and McCarthy 2002; Harrison et al. 2010; Leutner et al. 2012).

**Table 1.** Definitions of seven understory plant functional groups and predictions for how species richness of each group will vary with topography and microclimate in the 20.2 ha Niobrara Forest Dynamics Plot, North-Central Nebraska, USA. Plant functional groups were defined based on their structure, mature size, and physiological and morphological traits. Woody species includes woody plants and tree seedlings, and herbaceous species includes all other functional groups. C_3_ and C_4_ refer to the photosynthetic pathways.

| Functional group | Definition | Predictions for species richness |
| --- | --- | --- |
| Understory woody plants | Woody species that remain in the forest understory and/or retain a small stature across life stages (including lianas); broadleaved; C<sub>3</sub> photosynthetic pathway | Higher in moister, sheltered, shadier forested microrefugia and lower in the higher-light, more exposed floodplain and forest-prairie ecotone |
| Tree seedlings | Woody species (trees) that transition from the forest understory to the mid- and/or overstory across life stages; broadleaved; C<sub>3</sub> | Higher in forested microrefugia and lower in exposed floodplain and forest-prairie ecotone |
| Sedges | Non-woody, short-statured, perennial graminoids with triangular culms; narrow-leaved; C<sub>3</sub> | Higher outside of forested microrefugia in the exposed floodplain and forest-prairie ecotone |
| C<sub>3</sub> grasses | Non-woody, short-statured cool season graminoids; narrow-leaved; C<sub>3</sub> | Higher outside of forested microrefugia in the exposed floodplain and forest-prairie ecotone |
| C<sub>4</sub> grasses | Non-woody, short-statured warm season graminoids;<br>narrow-leaved; C<sub>4</sub> photosynthetic pathway | Higher outside of forested microrefugia in the<br>exposed floodplain and forest-prairie ecotone |
| Forbs | Non-woody, short-statured annual and perennial<br>flowering plants; broadleaved; C<sub>3</sub> | Higher in forested microrefugia and lower in<br>exposed floodplain and forest-prairie ecotone |
| Non-vascular and<br>lower vascular plants | Non-woody species that lack xylem and phloem<br>(bryophytes) and spore-bearing vascular plants (ferns,<br>horsetails, club/spike mosses); leaf structure variable; C<sub>3</sub> | Higher in forested microrefugia and lower in<br>exposed floodplain and forest-prairie ecotone |

We tested whether fine-scale microrefugia embedded within a broader regional climate-change refugium structure understory plant communities in a temperate forest. Understory plants are a key, but often understudied, component of forest diversity (Gilliam 2014; Landuyt et al. 2019; Spicer et al. 2022). Herbaceous plants alone comprise over 80% of vascular plant diversity in temperate forests (Gilliam 2007; Linares-Palomino et al. 2009; Spicer et al. 2020). Understory plants provide critical habitat and food resources for pollinators, specialist herbivores, and other faunal communities largely distinct from those supported by the overstory. As a result, declines in understory plant diversity can trigger cascading losses of dependent animal diversity (Masters et al. 1996; Hanula et al. 2015; Campbell et al. 2018; Cardoso et al. 2023). As climate change, land-use change, and forest fragmentation restructure forests worldwide (Hansen et al. 2013; Rodríguez-Echeverry et al. 2018; Albert et al. 2023), characterizing current patterns of understory plant diversity in wooded ecosystems is critical for identifying conservation priorities and maintaining forest biodiversity.

We hypothesized that spatial variation in microclimate buffering driven by topography, hydrogeology, and canopy structure generates localized microrefugia that shape understory plant diversity, composition, and distributions. Our study site is in a post-glacial migration corridor in the Niobrara River Valley, north-central Nebraska, USA. The warm, semi-arid macroclimate of the North American Great Plains largely supports temperate grassland (Whittaker 1975), but the Middle Niobrara River Valley has long been recognized as a regional climate-change refugium (Bessey 1887; Tolstead 1942a,b; Kaul et al. 1988). Seeps and springs fed by the High Plains aquifer (Ajaz et al. 2020) emerge along canyon walls draining into the Niobrara River, sustaining forests and supporting a distinctive assemblage of plant species from western, boreal, and eastern geographic provenances alongside tallgrass, mixed-grass, and sandhills prairie taxa (Bessey 1887; Hearty 1978; Churchill et al. 1988; Kantak 1995). This biogeographic crossroads – recognized as the Middle Niobrara River Valley Biologically Unique Landscape – supports many plant species found nowhere else in Nebraska or the broader region (Schneider et al. 2011). Within this regional refugium, fine-scale variation in topography, hydrogeology, and overstory structure co-determines microclimate, producing significantly cooler, moister, and less exposed conditions in lower-elevation forested canyons and upland forest than in the higher-elevation forest-prairie ecotone (McNichol et al. 2024). Despite the ecological significance of this refugium, understory woody and herbaceous plant distributions have not been surveyed for over three decades (Tolstead 1942a; Churchill et al. 1988; Kaul et al. 1988). Quantitative relationships between understory diversity, topography, and microclimate remain uncharacterized – a critical gap, given recent shifts in temperature and precipitation regimes and the accelerating pace of climate change globally (Diffenbaugh and Field 2013; IPCC 2022).

We collected data on the diversity, composition, and distributions of understory plant species across seven functional groups – woody plants, including tree seedlings, sedges, C_3_ and C_4_ grasses, forbs, and non-vascular and lower vascular plants (Table 1). We expected that microrefugia (as defined in Section 2.1) within the regional refugium would support higher diversity of mesic-adapted woody and herbaceous species, whereas the more exposed habitats along the floodplain and forest-prairie ecotone would favor drought-tolerant and light-demanding functional groups. Specifically, (P1) understory woody plant and tree seedling density and diversity and herbaceous plant richness should vary predictably with topography and microclimate, with functional group-specific predictions detailed in Table 1. We expected herbaceous functional groups, particularly forbs, to show weaker responses than woody plants, reflecting greater within-group variation in ecological strategies. (P2) Woody and herbaceous species should exhibit high turnover along topographic gradients, resulting in high beta diversity driven by shifts in species composition from microrefugia to the forest-prairie ecotone.

## 2. Methods

### 2.1 Study site, regional refugium, and microrefugia

The 20.2-ha Niobrara Forest Inventory Plot (hereafter, Niobrara Plot; 42°46’48.83” N, 100°01’15.56” W; Figure 1) is part of the Smithsonian Forest Global Earth Observatory (ForestGEO) network (Davies et al. 2021) and was established in 2019 within The Nature Conservancy’s Niobrara Valley Preserve (near Johnstown, Nebraska, USA; McNichol et al. 2024) in the Middle Niobrara River Valley. The Niobrara Plot (560 m × 360 m) is subdivided into 504 georeferenced 20 × 20 m quadrats (Figure 1c) and spans a sharp desiccation gradient driven by topography and hydrogeology from the Niobrara River to the Sandhills grassland (McNichol et al. 2024). McNichol et al. (2024) monitored microclimate variables at ten stations across this gradient from April-November in 2021 and 2022, documenting significant variation among five topographically defined habitats ranked from most mesic to most xeric: canyon bottoms, upland forest, upper canyon, floodplain, and ecotone. As the Niobrara Plot also samples the surrounding dominant Sandhills grassland, quadrats with zero woody stems ≥1 cm diameter at breast height (DBH; 1.3 m) were designated as grassland habitat (Figure 1b). To identify which habitats function as microrefugia – discrete areas within the regional refugium where local conditions are most strongly buffered from the regional macroclimate – we re-analyzed these data by calculating daily anomalies in maximum daily air temperature, maximum daily vapor pressure deficit (VPD), and minimum daily soil moisture at nine forested stations (canyon bottoms, upland forest, upper canyon, and floodplain) relative to an open prairie station representing ambient macroclimate conditions. Negative anomalies in maximum air temperature and VPD and positive anomalies in minimum soil moisture indicate microclimate buffering. Canyon bottoms, upland forest, and upper canyon consistently exhibited the largest buffering anomalies across both growing seasons and were designated as microrefugial habitats (see Figure S1 in Supporting Information). The floodplain showed partial buffering in soil moisture and VPD but positive temperature anomalies owing to its high-light, more exposed conditions, and was designated as outside of microrefugia.

**Figure 1.**
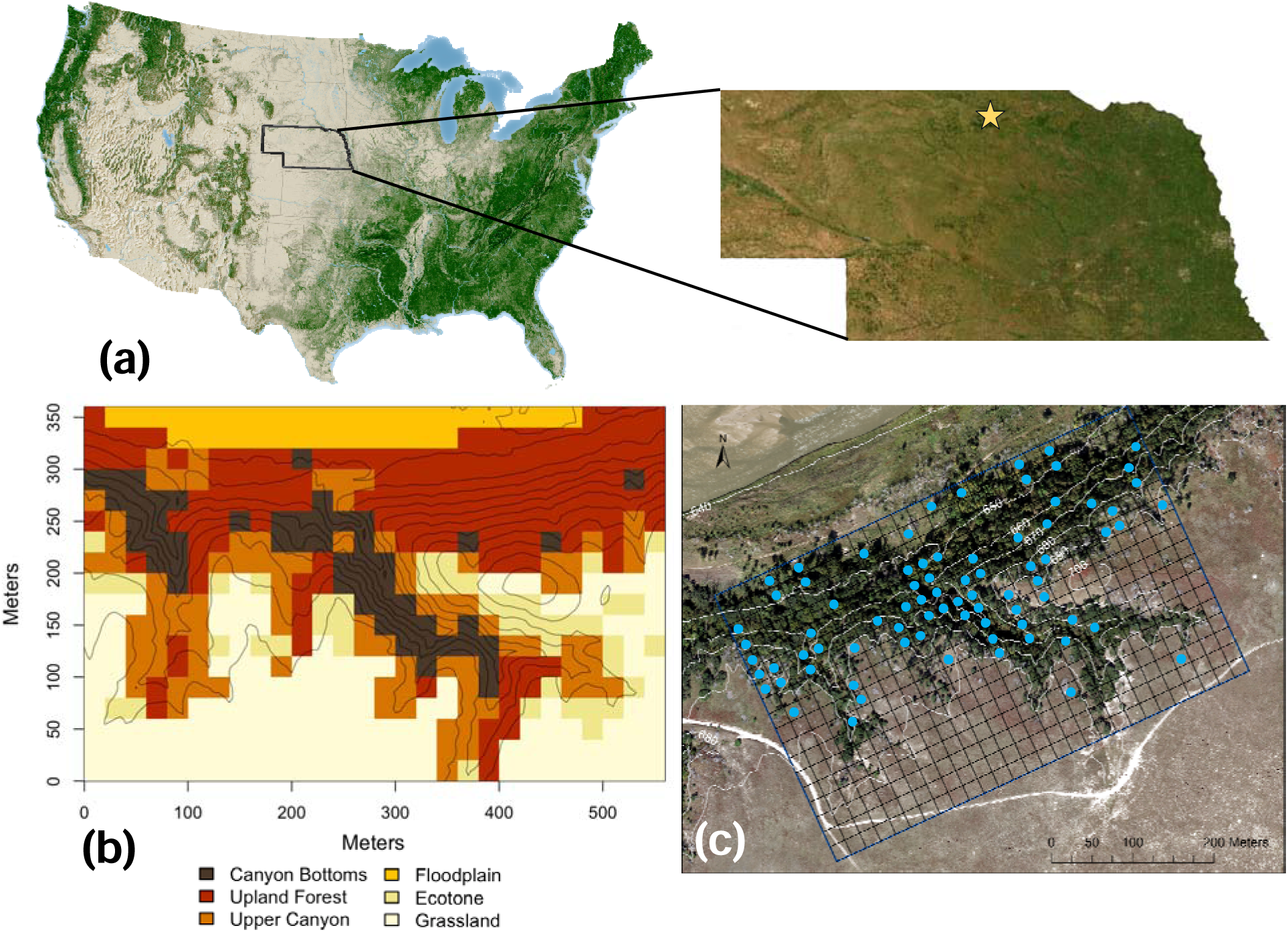
Geographic location, topographic habitats and understory plant sampling locations in the 20.2 ha Niobrara Forest Dynamics Plot, Niobrara River Valley, Nebraska, USA. (a) Location of the Niobrara Plot (yellow star) within Nebraska (15-m resolution aerial map; ESRI, 2022) and relative to forest cover across the continental United States (Forest Atlas of the United States, U.S. Forest Service). (b) Map of topographically defined habitats overlaid with 5-m elevation contour lines; habitat types (canyon bottoms, upland forest, upper canyon, floodplain, and ecotone) were defined in (McNichol et al. 2024); quadrats with zero woody stems with diameter at breast height 1 cm were designated as grassland. (c) High-resolution RGBN aerial image (UltraCam Eagle camera, Vexcel Imaging; 18 September 2019) showing the plot boundary, 20 × 20 m quadrats, and 10-m elevation contours (white lines). Understory woody plants were censused in a 1 × 1 m subplot at the northeastern corner of all 504 quadrats; herbaceous plants were censused in 78 subplots (blue points; canyon bottoms = 20; upland forest = 22; upper canyon = 18; floodplain = 9; ecotone = 9). Panels (a) and (c) modified from McNichol et al. 2024.

### 2.2 Woody and herbaceous plant censuses

In the initial 2019 census, all woody stems ≥1 cm DBH were tagged, mapped, identified to species, and measured for DBH, totaling 8,299 individuals of 27 species (McNichol et al. 2024). In May-July 2021, we established 504 permanently marked 1 × 1 m understory plant subplots at the northeastern corner of each quadrat and identified all woody stems <1 cm DBH to species; subplots were recensused in June-July 2022. In mid-July to early August 2021, we censused herbaceous plants in a stratified random subset of 78 subplots selected to represent the full range of topographic and microclimatic conditions within the forest and along the ecotone (Figure 1c). Subplots were distributed across habitat types as follows: canyon bottoms = 20; upland forest = 22; upper canyon = 18; floodplain = 9; ecotone = 9. Additional haphazard surveys targeting at-risk plant species were conducted across the plot during the 2023 tree recensus. In each subplot, we recorded presence of all forbs, graminoids, non-vascular, and lower vascular plants, and photographed and vouchered specimens that could not be field identified as morphospecies for later identification. Herbaceous plants were assigned to five functional groups: sedges, C_3_ grasses, C_4_ grasses, forbs, and non-vascular and lower vascular plants.

### 2.3 Estimation of topographic variables and measurements of soil moisture and light availability

Topographic variables were derived following McNichol et al. (2024). Briefly, a 1-m resolution Digital Elevation Model derived from LiDAR data (U.S. Geological Survey, 2017) was used to estimate the elevation (m), slope (%), northness, and eastness for each subplot. We quantified surface soil moisture by measuring volumetric water content (VWC, %) with a time-domain reflectometer (Hydrosense II, Campbell Scientific, Logan, UT USA) at three locations per subplot on rain-free days, six times between May and October 2021, across 73 subplots (canyon bottoms = 15; upland forest = 14; upper canyon = 14; floodplain = 15; ecotone = 15). We measured canopy openness (%) in July 2022 using a spherical concave densiometer at all subplots, averaging measurements in four cardinal directions. In a subset of 68 subplots with concurrent VWC measurements, we also took hemispherical canopy photographs 1 m above the soil surface before sunrise or after sunset using a Nikon CoolPix 4500 camera fitted with an FC-E8 fisheye lens, analyzing images in WinSCANOPY (Regent Instruments, Canada) to estimate canopy openness and mean daily photosynthetically active photon flux density (PPFD, mol/m^2^/day). Because densiometer-estimated openness was strongly correlated with hemispherical photo openness (*r* = 0.95) and PPFD (*r* = 0.93; Figure S2), only densiometer estimates were used in analyses.

### 2.4 Statistical analyses

Analyses were conducted in R 4.2.1 (R Core Team, 2022), using the ‘broom’, ‘car’, ‘dplyr’, ‘gghalves’, ‘ggplot2’, ‘MASS’, ‘mgcv’, ‘MuMIn’, ‘performance’, ‘pscl’, ‘RColorBrewer’, ‘tidyverse’, ‘tidyr’, ‘stats’, ‘vegan’, ‘viridis’, and ‘xlsx’ packages (Venables and Ripley 2002; Zeileis et al. 2008; Wood 2011; Wickham 2016; Fox and Weisburg 2019; Wickham et al. 2019; Lüdecke et al. 2021; Bartón 2022; Neuwirth 2022; Oksanen et al. 2022; R Core Team 2022; Wickham and Girlich 2022; Dragulescu and Arendt 2023; Robinson et al. 2023; Wickham et al. 2023a,b; Garnier et al. 2024; Tiedemann 2026). Analyses of all woody plants and tree seedlings were conducted separately for 2021 and 2022; subplots were the unit of replication, and statistical significance was assessed at = 0.05.

#### P1 Understory woody plant and tree seedling density and diversity and herbaceous plant richness vary with topography and microclimate

We calculated stem density, species richness, and Shannon’s diversity [*H* = Σ ln ; = proportion of individuals in the *i*th species] (Magurran 2004) for each subplot for (1) all understory woody species and (2) tree seedlings only (as species reaching the overstory may respond differently to topography and microclimate). Analyses included subplots with at least one woody stem (2021: *N* = 355; 2022: *N* = 346) or tree seedling (2021: *N* = 307; 2022: *N* = 306). For herbaceous plants (*N =* 78 subplots), we calculated species richness as a percent of the subplot total for each functional group, and the ratio of woody to non-woody species (number of woody species divided by the number of non-woody species, rWNW; *N* = 76, two subplots with zero non-woody species). Non-vascular and lower vascular plants were excluded from analyses due to low sample sizes (Table S1).

We evaluated topographic effects using two approaches. First, we used one-way analyses of variance (ANOVAs) to test for differences among habitat types (Figure 1b) in stem density, species richness, and Shannon’s diversity of understory woody plants and tree seedlings, percent richness of herbaceous functional groups, and rWNW. Response variables were transformed to meet model assumptions, and significant omnibus tests were followed by Tukey’s Honestly Significant Difference (HSD) comparisons with a 5% family-wise error rate.

Second, to account for collinearity among continuous topographic predictors, we conducted a principal components analysis (PCA) on the topographic variables (elevation, slope, northness, and eastness), retaining the first two PC axes as topographic predictors. Overstory basal area in a 10-m radius around each subplot was considered as a covariate but excluded due to collinearity with canopy openness (*r* = -0.83; Figure S3). We fit generalized additive models (GAMs) of woody plant and tree seedling richness and stem density with topographic PC1 and PC2 and openness as additive predictors. Prior to model fitting, we confirmed significant spatial autocorrelation in residuals using Moran’s I (all *p* < 0.001), justifying inclusion of a two-dimensional smooth of subplot coordinates to account for spatial structure (Hutchinson and Gessler 1994). GAMs were fit with Gaussian error distributions and identity link functions using penalized thin-plate regression splines (Hastie and Tibshirani 1987), with response variables transformed to meet model assumptions and fit assessed by *R*^2^.

For herbaceous functional groups and rWNW, we conducted the same PCA on topographic variables as described for woody plant models. We fit a single model per response variable with topographic PC1 and PC2 and openness as additive predictors; Moran’s I confirmed no significant spatial autocorrelation for any herbaceous functional group (all *p* > 0.05), so no spatial correction was applied. Exploratory analyses indicated zero inflation in richness of sedges, C_3_ and C_4_ grasses, and rWNW, so we fit zero-inflated negative binomial models for these response variables; for forbs we fit a binomial generalized linear model (GLM) with a logit link, modeling proportional richness with total subplot richness as weights. Model fits were assessed using McFadden’s pseudo-*R*² (p*R*²; McFadden 1974).

We tested whether understory plant species richness varied with soil moisture across the 42 herbaceous subplots with VWC measurements, conducting analyses separately for woody plants, tree seedlings, sedges, C_3_ grasses, C_4_ grasses, forbs, and rWNW. Mean and variation (standard deviation; SD) of VWC were calculated from the six measurements per subplot and standardized to zero mean and unit variance prior to analyses; mean and SD of VWC were not included in the same model due to collinearity (*r* = 0.64). For each response variable, we evaluated three candidate models: a null model, mean VWC with a quadratic term, and SD of VWC with a quadratic term. Quadratic terms were included to allow for unimodal moisture responses, and model support was evaluated using Akaike’s Information Criterion (AIC; Akaike, 1973) with Akaike weights. Moran’s I indicated no significant spatial autocorrelation in residuals for any functional group (all *p* > 0.05), so zero-inflated negative binomial models were fit without spatial correction for all response variables, including woody plants and tree seedlings, alongside a binomial GLM for forbs. Model fits were assessed using McFadden’s p*R*².

#### P2 Woody and herbaceous plants exhibit high beta diversity along topographic gradients

To visually assess species turnover, we plotted elevational distributions of understory woody species and herbaceous species with >1 individual as boxplots ordered by median elevation, with each stem (woody plants) or occurrence (herbaceous plants) assigned the elevation of its subplot. For woody plants, we quantified differences in species composition among habitats using permutational multivariate ANOVA (perMANOVA) with Cao dissimilarities as the response and habitat type as the predictor (Cao et al., 1997) and conducted a second perMANOVA using continuous topographic variables (elevation, slope, northness, eastness) as predictors. Scores from the first axis of a principal coordinates analysis (PCoA) of Cao dissimilarities were used to visualize woody plant compositional variation along topographic gradients. For herbaceous plants, we constructed a presence/absence community matrix across the 78 sampled subplots and calculated pairwise Sørensen dissimilarities (Bray-Curtis dissimilarity on presence/absence data). We conducted analogous perMANOVAs using Sørensen dissimilarities as the response, with habitat type and continuous topographic variables as predictors. To visualize herbaceous compositional variation along topographic gradients, we plotted scores from the first axis of a PCoA of Sørensen dissimilarities against subplot elevation.

## 3. Results

The 2021 understory woody plant census recorded 2,971 individuals of 30 species in 356 subplots (71%), and the 2022 recensus recorded 3,005 individuals of 30 species in 347 subplots (69%; Table S2). Across both years and the 2021 herbaceous plant census, we documented 170 total species: 14 understory woody plants, 17 tree seedlings, 7 sedges, 17 C_3_ grasses, 22 C_4_ grasses, 90 forbs, and 3 non-vascular and lower vascular plants (Figure 2a; Tables S1, S2). Across the 78 subplots censused for woody and herbaceous species, non-vascular and lower vascular plants and sedges were represented at low proportions in most subplots, C_3_ and C_4_ grasses at low proportions in several subplots, and understory woody plants, tree seedlings, and forbs at low proportions in few subplots (Figure 2b). Forbs were the most species-rich functional group across all habitats, followed by woody plants and tree seedlings in microrefugial habitats, and by C_3_ and C_4_ grasses in the exposed floodplain and ecotone (Figure 2c).

**Figure 2.**
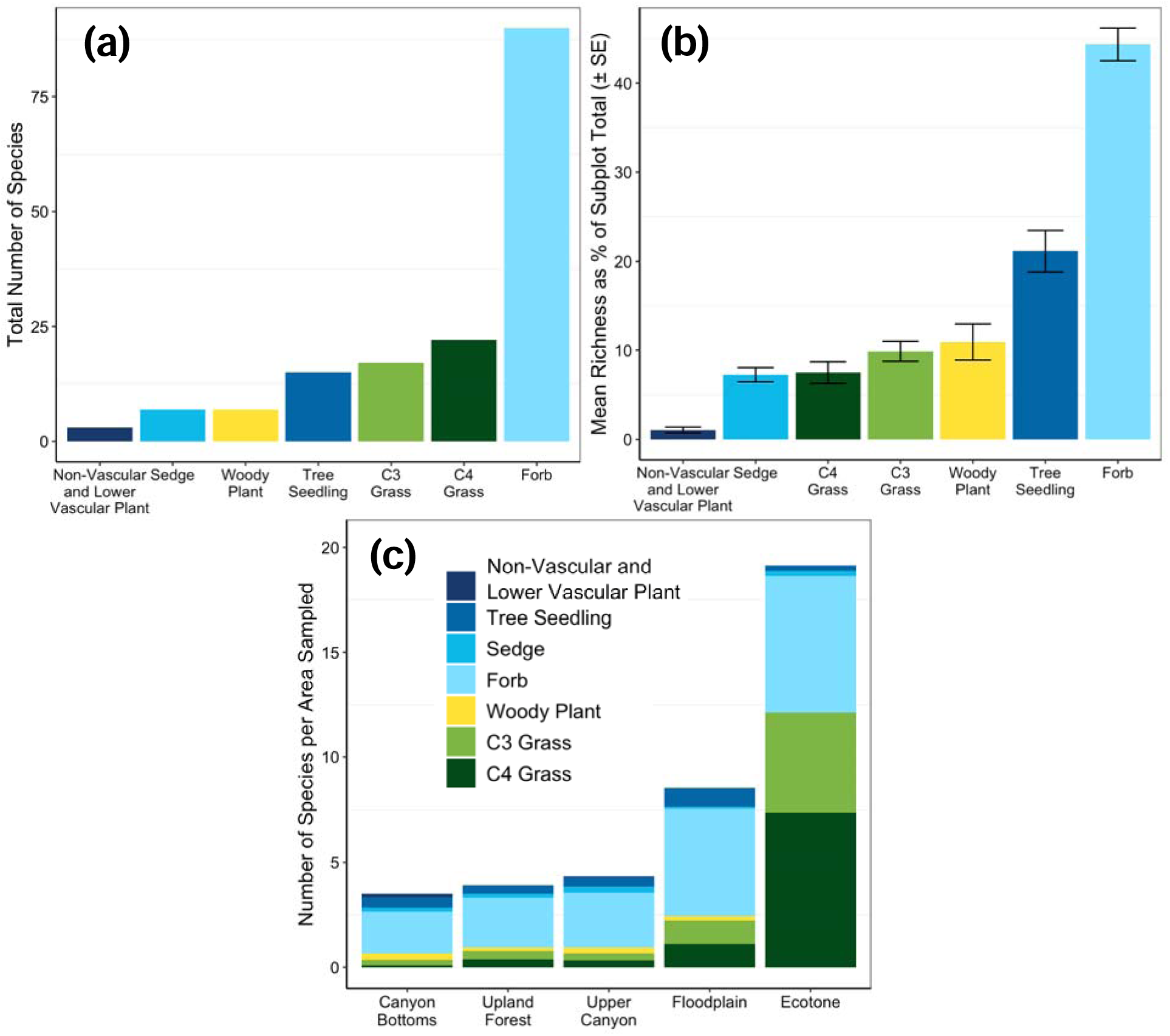
Patterns of understory woody and herbaceous plant species richness across 78 1 × 1 m subplots and among topographic habitats in the Niobrara Plot. (a) Total species richness per functional group; (b) mean (± standard error, SE) species richness per functional group as a percent of the subplot total; (c) total species richness per habitat standardized by area sampled, color-coded by functional group. The single grassland herbaceous subplot was grouped with the ecotone for all summaries and analyses. Herbaceous subplots sampled per habitat: canyon bottoms = 20; upland forest = 22; upper canyon = 18; floodplain = 9; ecotone = 9.

### 3.1 P1 Topography and microclimate effects on woody plant density and diversity and herbaceous plant richness

Understory woody species and tree seedling stem density and diversity varied among habitats in both years (Figures 3, S4; Table S3). Stem density was significantly higher in microrefugial habitats than in the ecotone and grassland (Figure 3a,b). Species richness and Shannon’s diversity showed a similar, but more pronounced pattern, with the highest values in the canyon bottoms and upland forest and lowest in the ecotone and grassland (Figures 3c-f). Tree seedlings showed analogous but weaker patterns (Table S3): stem density was only significantly higher in canyon bottoms and upland forest than grassland (Figure S4a,b) while richness and Shannon’s diversity were significantly higher in forested habitats than in both ecotone and grassland (Figure S4c-f).

**Figure 3.**
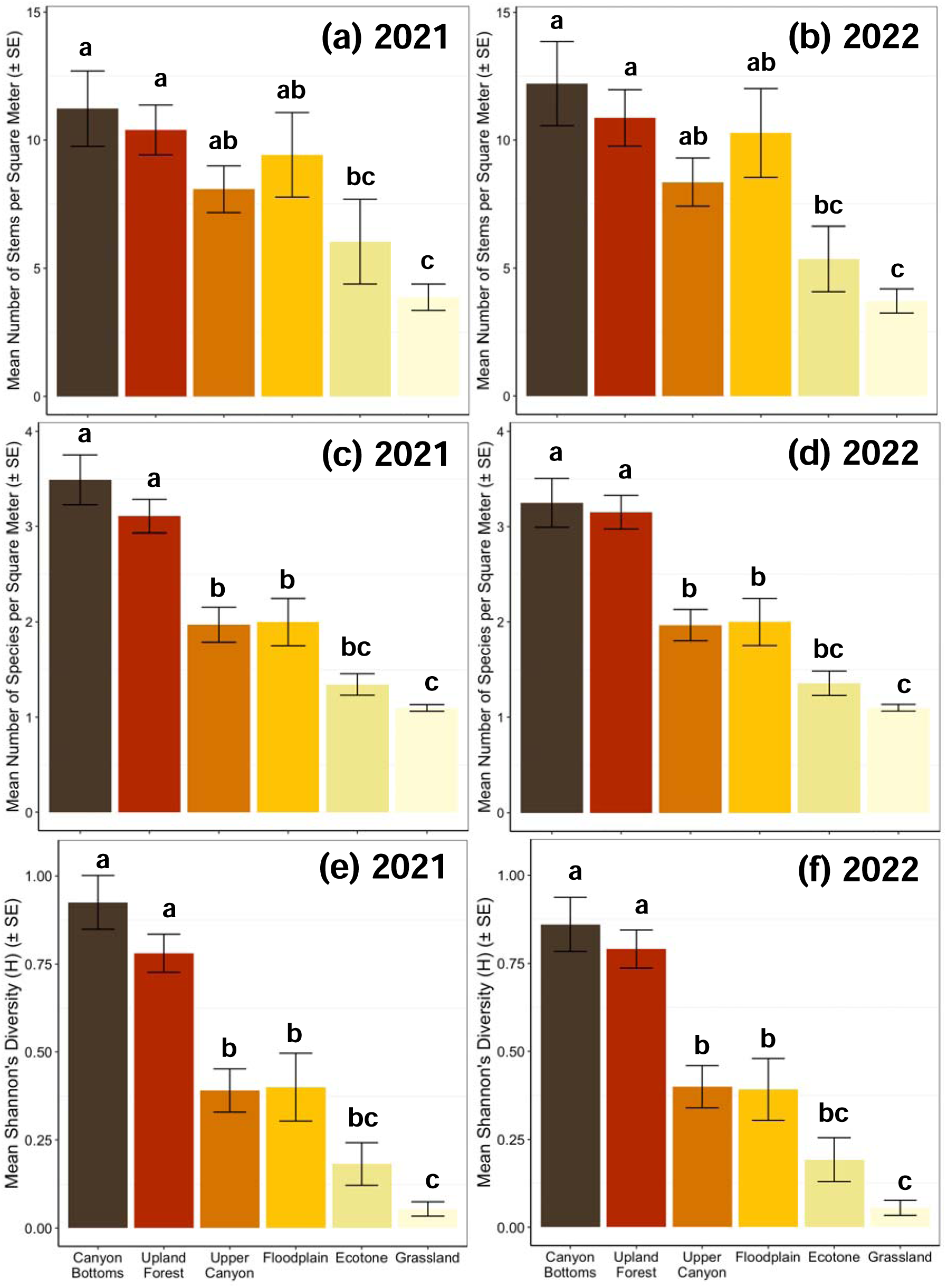
Variation in understory woody plant stem density, species richness, and Shannon’s Diversity among habitats in the Niobrara Plot. Habitat-related variation in mean (± standard error, SE) understory woody plant (diameter <1 cm) (a,b) stem density, (c,d) species richness, and (e,f) Shannon’s Diversity (H) per square meter in 2021 (a,c,e) and 2022 (b,d,f). Habitats are ordered by increasing solar radiation and exposure from canyon bottoms to grassland. Letters above the bars indicate significant pairwise differences at = 0.05 after correction for multiple comparisons using Tukey’s Honestly Significant Differences.

Herbaceous plant species richness and the ratio of the number of woody to non-woody species (rWNW) varied among habitats but patterns differed among functional groups (Figure 4; Table S4). C_3_ grass and forb richness did not differ significantly among habitats (Figure 4b,c), while sedge richness was significantly higher in the canyon bottoms than the floodplain (Figure 4a). C_4_ grass richness differed most strongly, with significantly higher richness outside of microrefugial habitats in the ecotone and floodplain (Figure 4d). Consistent with this, rWNW was significantly lower in the ecotone than the canyon bottoms (Figure 4e).

**Figure 4.**
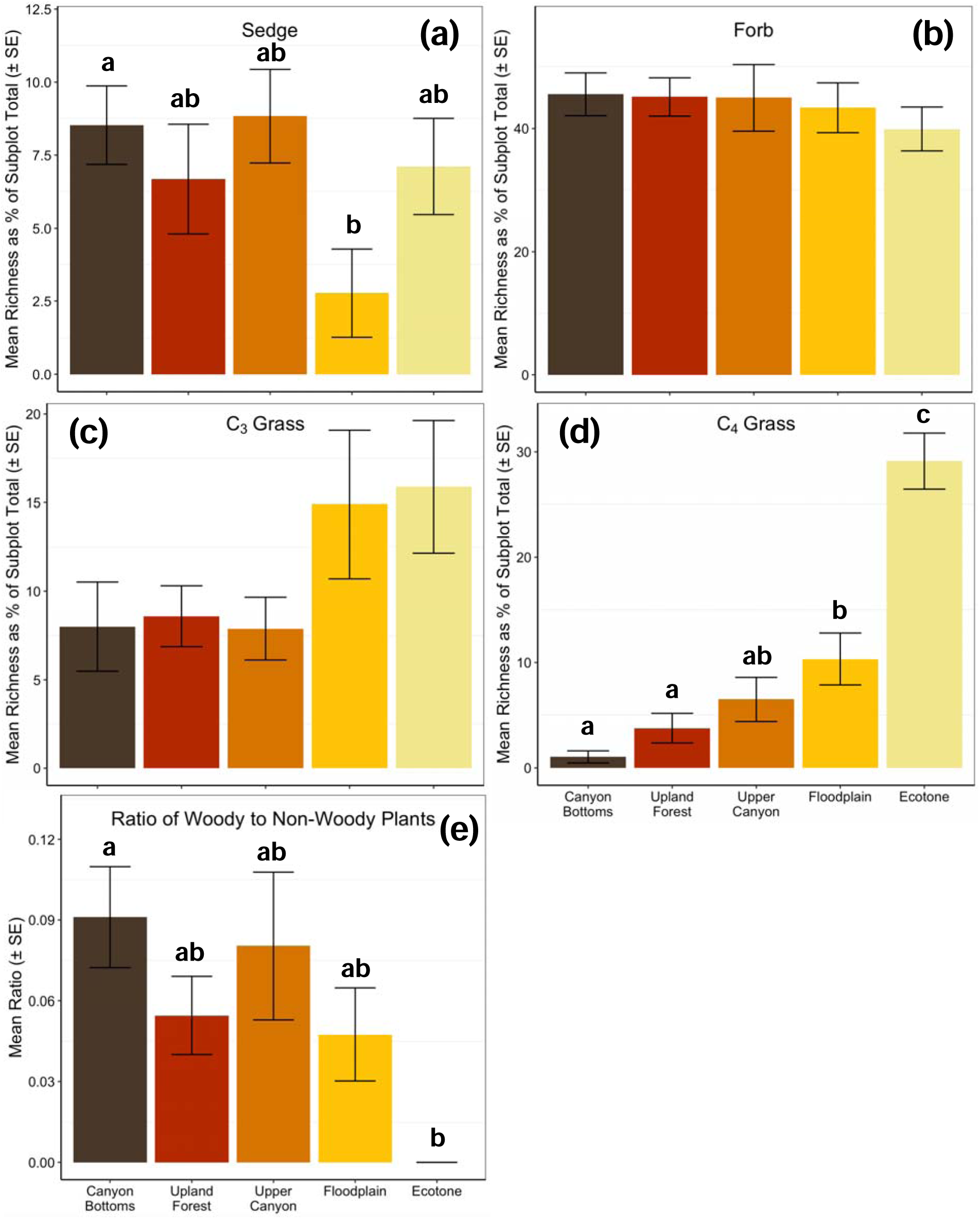
Variation in species richness of herbaceous functional groups and the ratio of woody to non-woody plants among habitats in the Niobrara Plot in 2021. Habitat-related variation in mean (± standard error, SE) species richness as a percent of subplot total for (a) sedges, (b) forbs, (c) C_3_ grasses, and (d) C_4_ grasses, and (e) the mean ratio of the number of woody to non-woody species. Habitats are ordered by increasing light intensity and exposure from canyon bottoms to ecotone. In (a,d,e), letters above the boxes indicate significant pairwise differences at = 0.05 after correction for multiple comparisons using Tukey’s Honestly Significant Differences.

Understory woody plant and tree seedling stem density and species richness varied significantly with canopy openness, with continuous topography playing a secondary role (Table S5). The first topographic PC axis (topoPC1; 30.8% of variation) contrasted lower-elevation, steep, north-facing positions characteristic of canyon microrefugia with higher-elevation, gentler, south-facing slopes, while the second axis (topoPC2; 25.6% of variation) captured east-west aspect variation. Stem density decreased significantly with increasing openness for both groups in both years (all *p* ≤ 0.01), while neither topographic PC axis was a significant predictor of stem density (Table S5). Species richness similarly declined significantly with increasing openness for woody plants and tree seedlings in both years (all p < 0.001; Table S5) and increased with higher topoPC1 values – indicating greater richness on lower-elevation, steep, north-facing slopes – in 2022 for woody plants (*p* = 0.033) and tree seedlings (*p* = 0.029). but not in 2021. TopoPC2 was not a significant predictor of richness in either year or group. Models explained substantially more variation in species richness (*R*² = 0.49–0.54) than in stem density (*R*² = 0.29–0.31).

Herbaceous species richness and rWNW showed variable responses to continuous topography and canopy openness, with weaker model fits than for woody plants (*pR*² = 0.021– 0.165; Table S6). C_4_ grasses showed the strongest responses: richness declined significantly with higher topoPC1 (*p* < 0.001), while the probability of absence declined with increasing topoPC1 (*p* = 0.029) and openness (*p* < 0.001), indicating that C_4_ grasses were both more likely to occur and more diverse outside of microrefugia. The rWNW declined significantly with higher topoPC1 (*p* = 0.045), suggesting that where woody and non-woody species co-occur, rWNW is lower in lower-elevation, steep, north-facing canyon positions. Sedges showed no significant effects of topography on richness but were more likely to be present in lower-elevation, north-facing, less exposed positions (topoPC1: *p* = 0.022; openness: *p* < 0.001). C_3_ grass richness increased marginally with openness (*p* = 0.040) but showed no significant topographic effects, nor did forbs.

Proportional species richness and rWNW also varied with soil moisture across functional groups, though effect sizes were generally modest (*pR*² = 0.033–0.115; Table 2; Figure S5). Mean VWC was the best-supported predictor for woody plants, C_3_ grasses, and rWNW, while the SD of VWC was best-supported for sedges, C_4_ grasses, and forbs. Woody plant richness showed significant quadratic relationships with mean VWC in both model components, and woody plants were significantly more likely to be present at lower mean VWC (all *p* < 0.05; Table 2). C_3_ grass richness declined significantly with mean VWC (*p* < 0.001) with a significant positive quadratic term (*p* = 0.009), indicating higher richness at both low and high moisture levels, and rWNW showed a similar but weaker pattern with significant effects only in the presence-absence component (Table 2). Sedge richness showed a significant positive quadratic relationship with the SD of VWC (*p* = 0.005), suggesting peak richness at intermediate moisture variability, while C_4_ grass richness declined significantly with the SD of VWC (*p* = 0.014). For tree seedlings, the null model received the most support (60.6% weight), with no significant effects of moisture on richness; however, tree seedling presence increased significantly at lower mean VWC (*p* = 0.037). Forbs showed no significant relationships with the SD of VWC.

**Table 2.** Top-supported models of understory woody and herbaceous plant species richness and the ratio of woody to non-woody species (rWNW) as a function of soil moisture in the Niobrara Plot in 2021. Candidate models included a null model, mean volumetric water content (VWC) with a quadratic term, and standard deviation of VWC (sdVWC) with a quadratic term; mean and SD VWC were modeled separately due to collinearity (*r* = 0.64). Analyses included the 42 subplots with soil moisture measurements. Parameter estimates, standard errors (SE), *z*-values, and *p*-values are shown separately for the count component and the zero-inflation component of zero-inflated negative binomial models; McFadden’s pseudo-*R*² (p*R*²) and Akaike weights are reported. VWC² and sdVWC² denote quadratic terms; significant effects at _α_ = 0.05 are shown in bold. *Forbs were modeled using a binomial generalized linear model with a single component; presence/absence columns not applicable (—).

| Functional group | $pR^2$ | Weight (%) | Predictor | Count component (richness \| present) | | | | Presence/absence component | | | |
| --- | --- | --- | --- | --- | --- | --- | --- | --- | --- | --- | --- |
| | | | | Estimate | SE | z-value | $p$-value | Estimate | SE | z-value | $p$-value |
| Woody plants | 0.079 | 94.9 | Intercept | 2.250 | 0.081 | 27.831 | <**0.001** | -0.843 | 0.507 | -1.664 | 0.096 |
|  |  |  | VWC | -0.289 | 0.141 | -2.050 | **0.040** | -2.215 | 0.833 | -2.660 | **0.008** |
|  |  |  | VWC<sup>2</sup> | 0.104 | 0.052 | 2.001 | **0.045** | 0.668 | 0.319 | 2.093 | **0.036** |
| Tree seedlings | 0.033 | 37.9 | Intercept | 2.748 | 0.120 | 22.898 | <**0.001** | -1.081 | 0.512 | -2.112 | **0.035** |
|  |  |  | VWC | 0.191 | 0.201 | 0.951 | 0.341 | -1.594 | 0.764 | -2.087 | **0.037** |
|  |  |  | VWC<sup>2</sup> | -0.070 | 0.117 | -0.593 | 0.553 | 0.764 | 0.371 | 2.061 | **0.039** |
| Sedges | 0.044 | 59.8 | Intercept | 2.254 | 0.115 | 19.574 | **<0.001** | -0.965 | 0.448 | -2.151 | **0.031** |
|  |  |  | sdVWC | -0.218 | 0.130 | -1.672 | 0.094 | -0.657 | 0.471 | -1.395 | 0.162 |
|  |  |  | sdVWC<sup>2</sup> | 0.250 | 0.089 | 2.804 | **0.005** | 0.347 | 0.271 | 1.281 | 0.200 |
| C<sub>3</sub> grasses | 0.037 | 55.8 | Intercept | 2.600 | 0.118 | 22.077 | **<0.001** | -1.241 | 0.480 | -2.583 | **0.010** |
|  |  |  | VWC | -0.612 | 0.182 | -3.367 | **<0.001** | 0.311 | 0.751 | 0.414 | 0.679 |
|  |  |  | VWC<sup>2</sup> | 0.195 | 0.074 | 2.622 | **0.009** | -0.073 | 0.313 | -0.233 | 0.816 |
| C<sub>4</sub> grasses | 0.040 | 42.7 | Intercept | 3.039 | 0.125 | 24.286 | **<0.001** | 0.060 | 0.404 | 0.149 | 0.882 |
|  |  |  | sdVWC | 0.026 | 0.135 | 0.195 | 0.846 | 0.597 | 0.463 | 1.287 | 0.198 |
|  |  |  | sdVWC<sup>2</sup> | -0.190 | 0.077 | -2.468 | **0.014** | -0.263 | 0.267 | -0.988 | 0.323 |
| rWNW | 0.064 | 87.7 | Intercept | 2.500 | 0.090 | 27.845 | **<0.001** | -0.843 | 0.507 | -1.664 | 0.096 |
|  |  |  | VWC | -0.236 | 0.162 | -1.454 | 0.146 | -2.238 | 0.842 | -2.659 | **0.008** |
|  |  |  | VWC<sup>2</sup> | 0.076 | 0.060 | 1.269 | 0.204 | 0.682 | 0.326 | 2.093 | **0.036** |
| Forbs* | 0.115 | 42.3 | Intercept | 0.214 | 0.109 | 1.970 | **0.049** | — | — | — | — |
|  |  |  | sdVWC | 0.108 | 0.126 | 0.856 | 0.392 | — | — | — | — |
|  |  |  | sdVWC<sup>2</sup> | 0.044 | 0.070 | 0.626 | 0.532 | — | — | — | — |

### 3.2 P2 Topography generated high species turnover and beta diversity for understory woody and herbaceous plants

Topography strongly structured understory woody species distributions, with species turnover generating high beta diversity both years (Figures 5, S6; Table S7). Woody species occupied distinct elevation niches (Figures 5a, S6a); some were restricted to narrow elevational ranges, indicating strong habitat affinities (*e.g.*, *Salix* spp., *Rosa arkansana*), while others spanned broader gradients (*e.g.*, *Celtis occidentalis*, *Prunus americana*, *Rubus occidentalis*). Alpha diversity was highest in forested habitats (Figures 5b, S6b). Species composition differed significantly among habitats, explaining 21.6% and 24.0% of variation in 2021 and 2022, and varied with continuous topography (Figures 5c, S6c), with elevation and slope explaining more variation (6-17%) than eastness and northness (<1%; Table S7).

**Figure 5.**
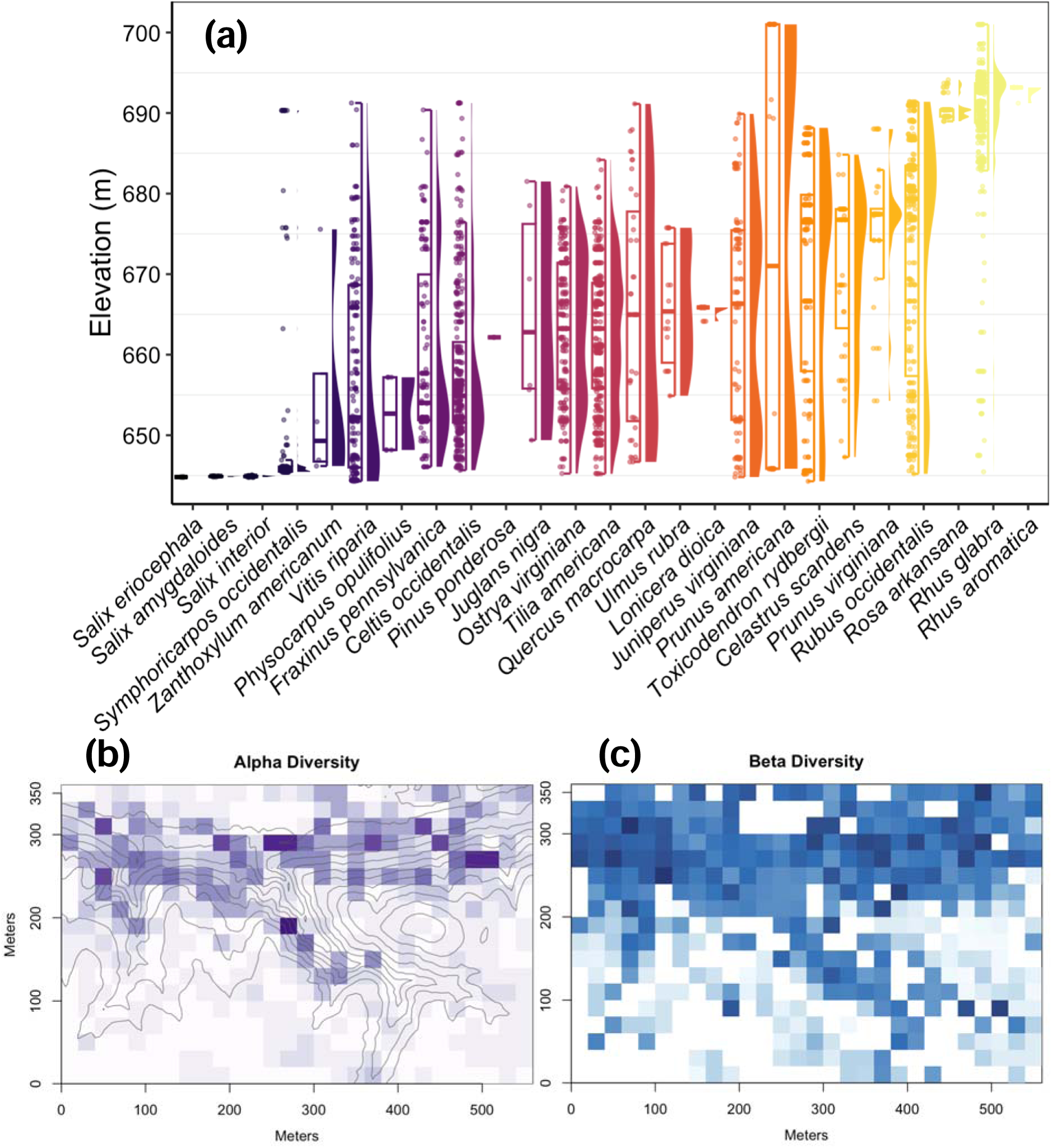
Understory woody plant species turnover and alpha and beta diversity with respect to elevation in the Niobrara Plot in 2021. (a) Combined box and violin plots showing the elevation distributions for woody species with >1 individual, ordered by increasing median elevation. Boxes indicate the median (center line) and first and third quartiles; lower and upper whiskers indicate the first and third quartiles ±1.5 times the interquartile range, and points 51 indicate outliers; (b) heatmap showing species richness (alpha diversity) of 1 × 1 m subplots overlaid with 5-m elevation contour lines; 20 × 20 m quadrats are shaded from light to dark purple with increasing woody species richness (from 0-9 species) in the nested subplot; (c) heatmap showing variation in woody species composition (beta diversity) overlaid with 5-m elevation contour lines; quadrats are color-coded by scores from the first axis of a principal coordinates analysis of Cao dissimilarities in subplot-level species composition, with similarly shaded squares indicating more similar species assemblages. In (b) and (c), white quadrats represent nested subplots containing no woody stems <1 cm diameter at breast height.

Herbaceous species distributions also varied with topography, though less strongly and consistently than woody plants (Figures 6, S7; Table S8). Graminoids occupied distinct topographic niches, with several C_3_ and C_4_ grasses concentrated at lower elevations in the exposed floodplain (*e.g.*, *Spartina pectinata*) and higher elevations along the ecotone (*e.g.*, *Bouteloua hirsuta*, *Dichanthelium wilcoxianum*), while other grasses and sedges had broader elevational distributions (Figure 6a). Non-vascular and lower vascular plants were concentrated in mid-elevation canyon microrefugia, whereas forbs spanned the full elevational range, including widespread generalists (*e.g.*, *Erigeron philadelphicus*, *Hackelia virginiana*) and specialists at lower (*e.g.*, *Boehmeria cylindrica*, *Mentha canadensis*) and higher elevations (*e.g.*, *Helianthus petiolaris*, *Lithospermum incisum*; Figure 6b). Herbaceous alpha diversity was highest in the floodplain and ecotone (Figure S7a). Species composition differed significantly among habitats, explaining 17.9% of variation (Table S8), and as with woody plants, slope and elevation explained more compositional variation (8.9% and 6.6%, respectively) than northness (2.5%) or eastness (not significant; Table S8). PCoA axis 1 explained 19.8% of variation in herbaceous composition and separated subplots along the elevational and exposure gradient; floodplain and ecotone subplots clustered together, and microrefugial habitat subplots formed a distinct group (Figure 6c).

**Figure 6.**
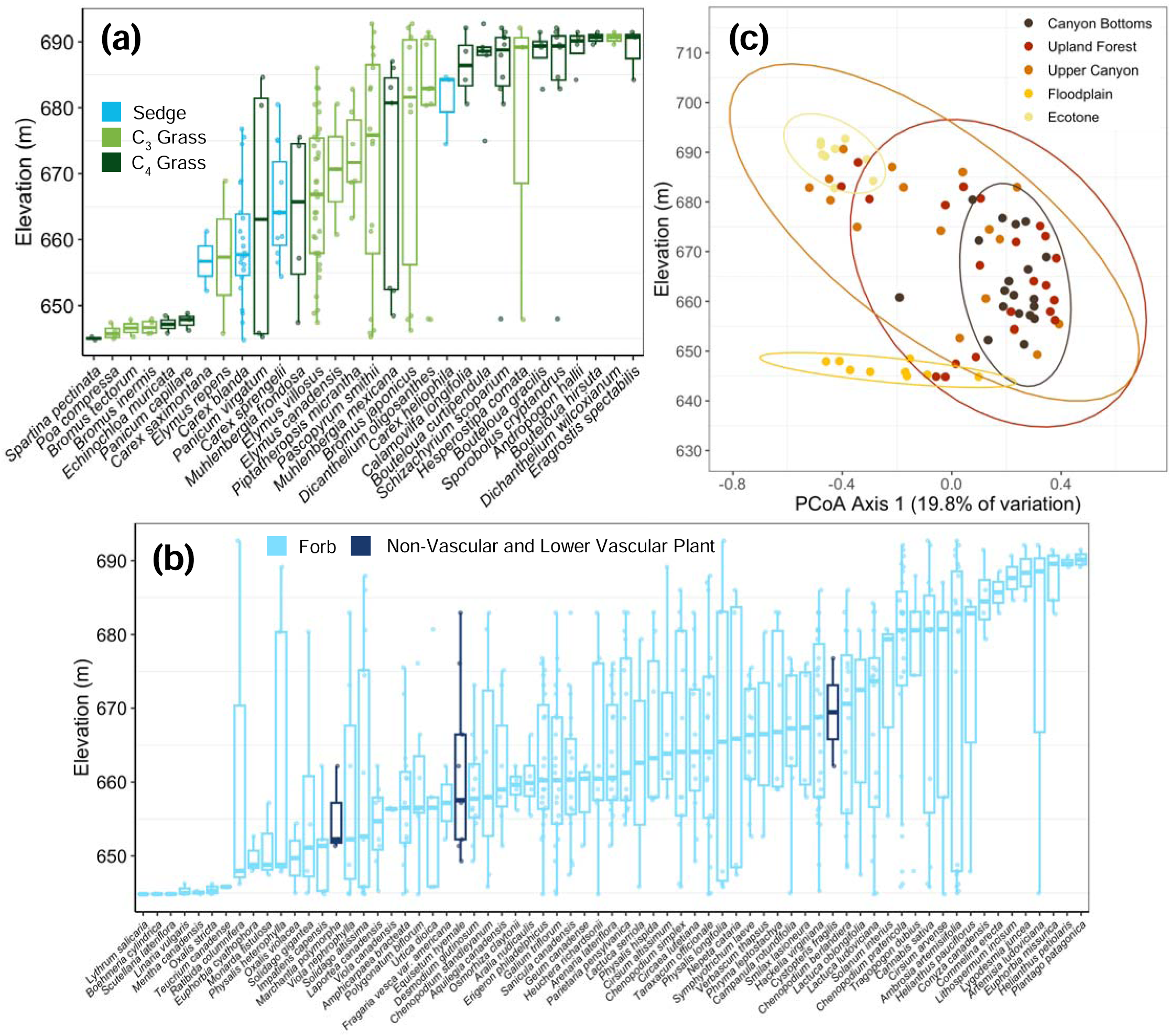
Herbaceous plant species turnover and compositional variation with respect to elevation in the Niobrara Plot in 2021. Boxplots showing the elevational distributions of (a) graminoid species and (b) forb and non-vascular and lower vascular plant species with >1 occurrence, ordered by increasing median elevation and color-coded by functional group ((a) C_3_ grasses: light green; C_4_ grasses: dark green; sedges: cyan; (b) forbs: light blue; non-vascular and lower vascular plants: navy). Boxes indicate the median (center line) and first and third quartiles; lower and upper whiskers indicate the first and third quartiles ± 1.5 times the interquartile range; points show individual subplot occurrences. (c) first axis scores from a principal coordinates analysis of Sørensen dissimilarities plotted against subplot elevation, with points color-coded by habitat type; ellipses represent 95% confidence regions. Species names shortened for visualization: (a) *Dicanthelium oligosanthes* var. *scribnerianum* to *D. oligosanthes*; (b) *Helianthus pauciflorus* subsp*. Subrhomboideus* to *H. pauciflorus*.

## 4. Discussion

As climate change accelerates and forested biomes become increasingly fragmented, major redistributions of forest biodiversity are expected (Hansen et al. 2013; Rodríguez-Echeverry et al. 2018; Albert et al. 2023), yet most understory plant communities in woody ecosystems remain understudied (Spicer et al. 2022). Here, we address this gap by quantifying woody and herbaceous plant diversity within a biogeographically important climate-change refugium sheltering woodland flora in the grassland that largely covers the semi-arid North American Great Plains. A novel contribution of our study is the direct measurement of multiple dimensions of microclimate alongside plant diversity to identify how understory diversity is structured by topographically driven microclimate gradients within the regional refugium. Consistent with patterns in the overstory (McNichol et al. 2024), topography and microclimate strongly structured understory plant diversity, composition, and distributions. Moister, shadier, less exposed microrefugia supported higher understory woody plant density and diversity, whereas C_4_ grass richness was highest along the exposed forest-prairie ecotone where microclimate buffering was weakest. These contrasting responses of functional groups to microclimate buffering generate strong species turnover within this regional refugium. Sedges and C_3_ grasses showed weaker habitat-level differentiation but responded to continuous variation in soil moisture – with sedge richness peaking at intermediate moisture variability and C_3_ grass richness higher at both moisture extremes – reflecting variation in species’ soil moisture affinities within these functional groups. Forb richness showed no significant responses to topography or microclimate, likely because this functionally diverse group includes species spanning the full range of drought and heat tolerance, from canyon microrefugia specialists to ecotone specialists, dampening any group-level environmental signal. Because regional refugia such as the Middle Niobrara River Valley have been critical in maintaining regional forest diversity through past climatic transitions (Bennett et al. 1991; Petit et al. 2003), our findings underscore the conservation importance of the Niobrara forests, where topographic complexity supports high beta diversity and increases regional gamma diversity.

### 4.1 Microrefugia structure understory plant diversity, composition, and species turnover

Woody species commonly exhibit specialization across topographic habitats (Kenfack et al. 2014; Zuleta et al. 2020). In the Niobrara forests, understory woody plant density and diversity were consistently lower in exposed ecotone and grassland than in microrefugial habitats with greater canopy cover, indicating strong habitat filtering. The degree of microclimate buffering – quantified by negative anomalies in maximum air temperature and VPD and positive anomalies in minimum soil moisture of forested areas relative to the prairie – also strongly structured woody plant density and diversity along continuous gradients, consistent with patterns in other forests (Nagamatsu et al. 2002; Muñoz-Rojas et al. 2016; Jin et al. 2018). Increased canopy openness was associated with reduced woody stem density and species richness, likely reflecting increasing desiccation stress and loss of canopy-mediated buffering, which can limit seedling survival (Gómez-Aparicio et al. 2008; Villegas et al. 2010; Davis et al. 2019). In contrast to studies reporting lower seedling survival on steep slopes (Nagamatsu et al. 2002; Abiem et al. 2023), woody seedling density and richness were higher on steeper slopes, where groundwater springs and seeps, cooler microclimates, and greater overstory complexity in the Niobrara canyons coincide to amplify buffering and generate microrefugia (Cartwright et al. 2020; McNichol et al. 2024; Button and Piovia-Scott 2025). The decline in richness at the highest soil moisture levels likely reflects inundation constraints in the floodplain (Glenz et al. 2006). Together, these patterns indicate that woody diversity is structured by fine-scale thermal, atmospheric, and hydrological variation.

Both woody plants and tree seedlings also showed strong niche partitioning and species turnover across the short elevational gradient. Seedlings of smaller-statured trees occurred in the low-elevation floodplain (*e.g.*, *Salix interior*), eastern deciduous canopy trees were concentrated in mid-elevation canyon bottoms and upland forest (*e.g.*, *Tilia americana*), and understory trees and shrubs occurred more frequently in the high-elevation ecotone and grassland (*e.g.*, *Prunus virginiana*). These results parallel topographic niche partitioning in the overstory of the Niobrara Plot (McNichol et al. 2024) and resemble topography-driven beta diversity in tropical forests across similarly short elevational distances (Harms et al. 2001; Pulla et al. 2017; Zuleta et al. 2020). This elevational partitioning translated into strong differences in species composition among habitats. Eight woody species — including *Betula papyrifera*, *Juglans nigra*, and *Pinus ponderosa* from northern, eastern, and western geographic provenances, respectively — were documented exclusively in microrefugial habitats both years. These findings indicate that although microrefugia occupy a relatively small portion of the landscape, they support distinct woody assemblages associated with cooler, less exposed, and more hydrologically buffered conditions.

Herbaceous functional groups showed less consistent habitat-level responses in species richness than woody plants, with significant effects on richness detected only for C_4_ grasses and rWNW. This likely results from high within-group variation in ecological strategies, which can dampen habitat-level signals when species with contrasting environmental affinities are aggregated (Pausas and Austin 2001). Nevertheless, herbaceous species showed strong but more variable elevational structuring and species turnover. Graminoids exhibited niche partitioning, with several C_3_ grasses concentrated at lower elevations in the exposed floodplain and higher elevations along the ecotone, and most sedges occurring at mid-elevations within microrefugia. Forbs spanned the full elevational range, while non-vascular and lower vascular plants were concentrated in mid-elevation canyon microrefugia. C_4_ grasses were primarily distributed in high-light, drier environments outside of microrefugia, consistent with physiological adaptations to heat and water stress, including high water-use efficiency, osmotic adjustments sustaining photosynthesis during drought (Knapp 1985; Monson et al. 2025), and utilization of transient shallow soil moisture (Eggemeyer et al. 2006). Among sedges, C_3_ grasses, and forbs, topography and canopy openness explained little variation in richness, although sedges were more likely to be present in north-facing, less exposed positions consistent with microrefugia, and C_3_ grass richness increased marginally with openness. This suggests that topography structures herbaceous richness primarily through fine-scale soil moisture gradients (Bennie et al. 2008; Rita et al. 2021), with patterns further reflecting biogeographic affinities (Churchill et al. 1988; Kaul et al. 1988).

Despite relatively weak functional group-level responses in richness, microrefugia harbored a significant concentration of individual herbaceous species, with all three non-vascular and lower vascular plant species (100%), five sedge species (71%), 30 forb species (33%), four C_3_ grass species (23%), and two C_4_ grass species (9%) found only in microrefugial habitats. Numerous herbaceous associates of North American eastern deciduous forests, including *Aralia nudicaulis* (Araliaceae), *Geum canadense* (Rosaceae), and *Osmorhiza claytonii* (Apiaceae) were restricted entirely to canyon bottom and upland forest microrefugia, consistent with their associations with cool, moist conditions across their more easterly and northerly geographic ranges. Opportunistic observations undertaken to supplement our systematic sampling documented additional species of regional conservation concern in microrefugial habitats, including the orchids *Liparis loeselii* (Orchidaceae; Nebraska S1: critically imperiled) and *Platanthera aquilonis* (Orchidaceae; Nebraska S3: vulnerable to extirpation), the mycoheterotroph *Pterospora andromedea* (Ericaceae; Nebraska S2: imperiled), and *Asclepias lanuginose* (Apocynaceae), a milkweed with range-wide declining populations. The concentration of species spanning multiple functional groups, including several of conservation concern, exclusively within microrefugial habitats suggests that habitat-level richness alone may underestimate their importance for maintaining regional plant diversity.

The semi-arid macroclimate of the region already limits woody expansion beyond the ecotone (Kaul et al. 1988; Muñoz-Rojas et al. 2016), while less frequent but more intense precipitation events may further alter water availability for plants (Singh et al. 2013; Bishop et al. 2021). However, persistence of the Niobrara microrefugia depends on more than precipitation: canopy-mediated reductions in temperature and VPD, increased soil moisture, complex overstory structure, and groundwater springs and seeps all contribute to the buffered conditions that distinguish these habitats from the surrounding prairie. Changes in precipitation regimes, warming, or hydrological connectivity could therefore erode different components of this buffering and alter the distributions of species currently restricted to microrefugial habitats.

These results highlight the importance of tracking changes in understory plant diversity and distributions along this fine-scale environmental gradient as regional climate conditions continue to change.

### 4.2 Conservation implications

Our results show that fine-scale environmental heterogeneity within regional climate-change refugia is particularly important in biogeographically complex regions where species with contrasting climatic affinities occur in close proximity. In the Middle Niobrara River Valley, topographic complexity, canopy structure, and groundwater inputs generate a mosaic of buffered and exposed microclimates that supports distinct understory assemblages and high species turnover across short spatial distances. Such microrefugia may help decouple local species assemblages from regional macroclimatic change by moderating heat and moisture stress that would otherwise exceed species’ physiological tolerances (Spector 2002; Hylander et al. 2015; McLaughlin et al. 2017; Davis et al. 2019; Morelli et al. 2020). This buffering may be particularly important where species from different biogeographic provenances persist near the edges of their geographic ranges (Slatyer et al. 2013; Sheth and Angert 2014), including boreal taxa restricted to cool, moist microsites (Stralberg et al. 2020a; McNichol et al. 2024). Our findings therefore suggest that conservation of regional climate-change refugia should account not only for their extent, but also for the topographic, canopy, and hydrological heterogeneity that generates and sustains microrefugia. Maintaining these features may help preserve climatically sensitive plant species and the distinct understory assemblages they support as regional warming and drying intensify, emphasizing the value of incorporating fine-scale environmental conditions into conservation planning for refugial landscapes (Stralberg et al. 2020b; Dobrowski et al. 2021).

## Acknowledgements

We thank Emma Decker and Susana Moyer for assistance with seedling censuses and Sakia Fields, Jennifer Foster, Joseph Eastman, Townson Lemansky, Jack Luth, Susana Moyer, Ashley Newman, Cooper Steffen, Patrick Wilson, and Jessica Shue for assistance with the first census of the Niobrara Plot. We acknowledge The Nature Conservancy for permission to install the Niobrara Plot on the Niobrara Valley Preserve and for providing logistical support. This manuscript is a substantially revised and expanded version of Chapter 2 of the first author’s doctoral dissertation (McNichol BH (2023) Effects of topography and microclimate on plant community structure in a Great Plains refugial forest. PhD dissertation, University of Nebraska– Lincoln, ProQuest Order No. 30569054). This research was supported by funding from Jacqueline B. Mars, the Smithsonian ForestGEO program, the National Science Foundation AccelNet Program (OISE 2020424), the American Philosophical Society, the Botanical Society of America, the Nebraska Native Plant Society, the University of Nebraska–Lincoln (UNL) Center for Plant Science Innovation, the UNL School of Biological Sciences Dr. John F. Davidson Memorial Fund, a UNL Faculty Seed Grant, the UNL Undergraduate Creative Activities and Research Experience program, and the Cabela’s Apprenticeship Program.

## Data Availability Statement

Data used in this manuscript are public on the Stewarding Nebraska Data (SANDY) research data repository at the University of Nebraska– Lincoln (https://sandy.unl.edu).

