## Supplementary material for "Environmental correlates of understory plant diversity and distribution in a temperate climate-change refugium": Seedling_SI_Final_DD.docx

**Title**

**Running Title**

Understory diversity in forest refugia

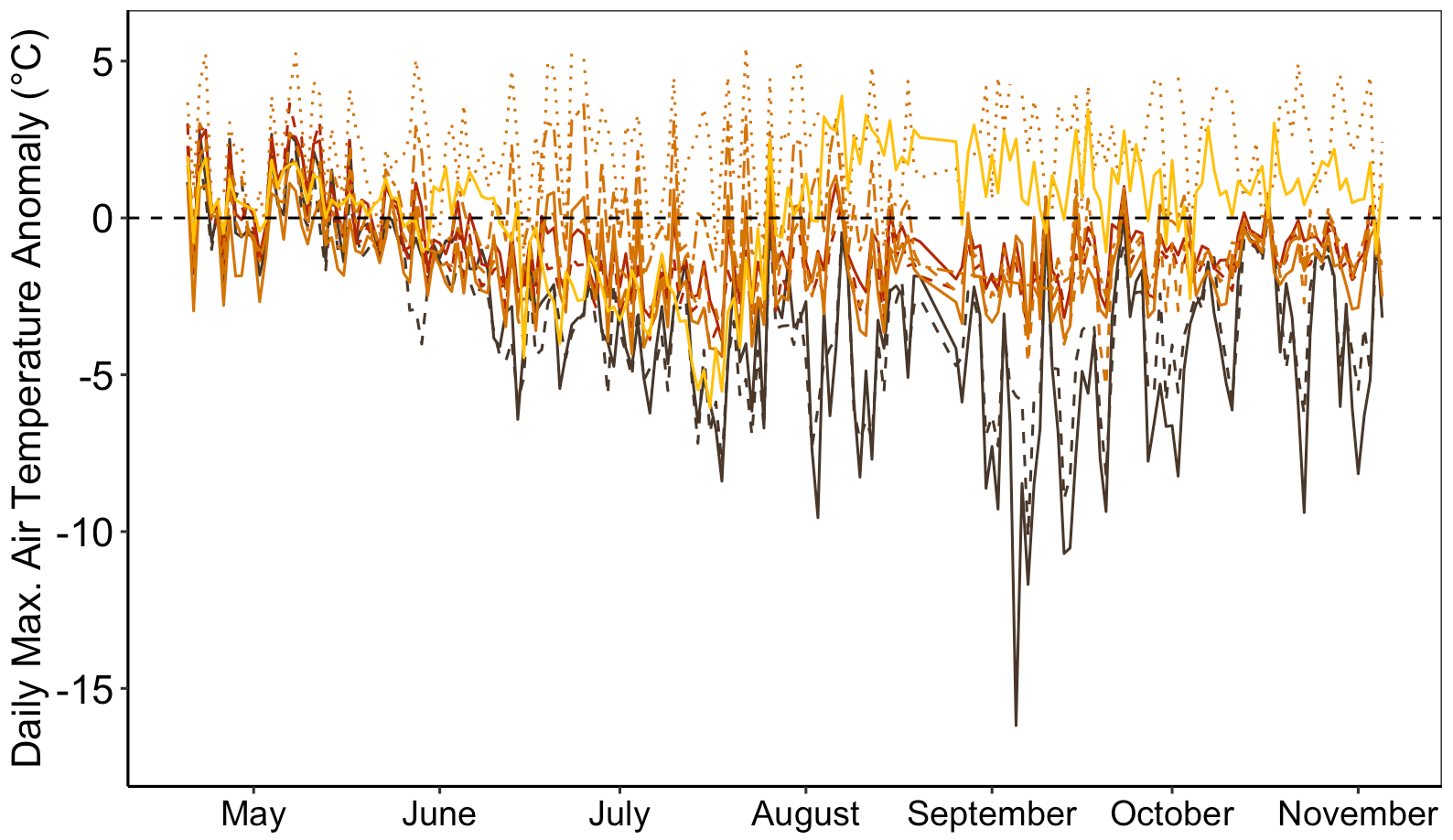

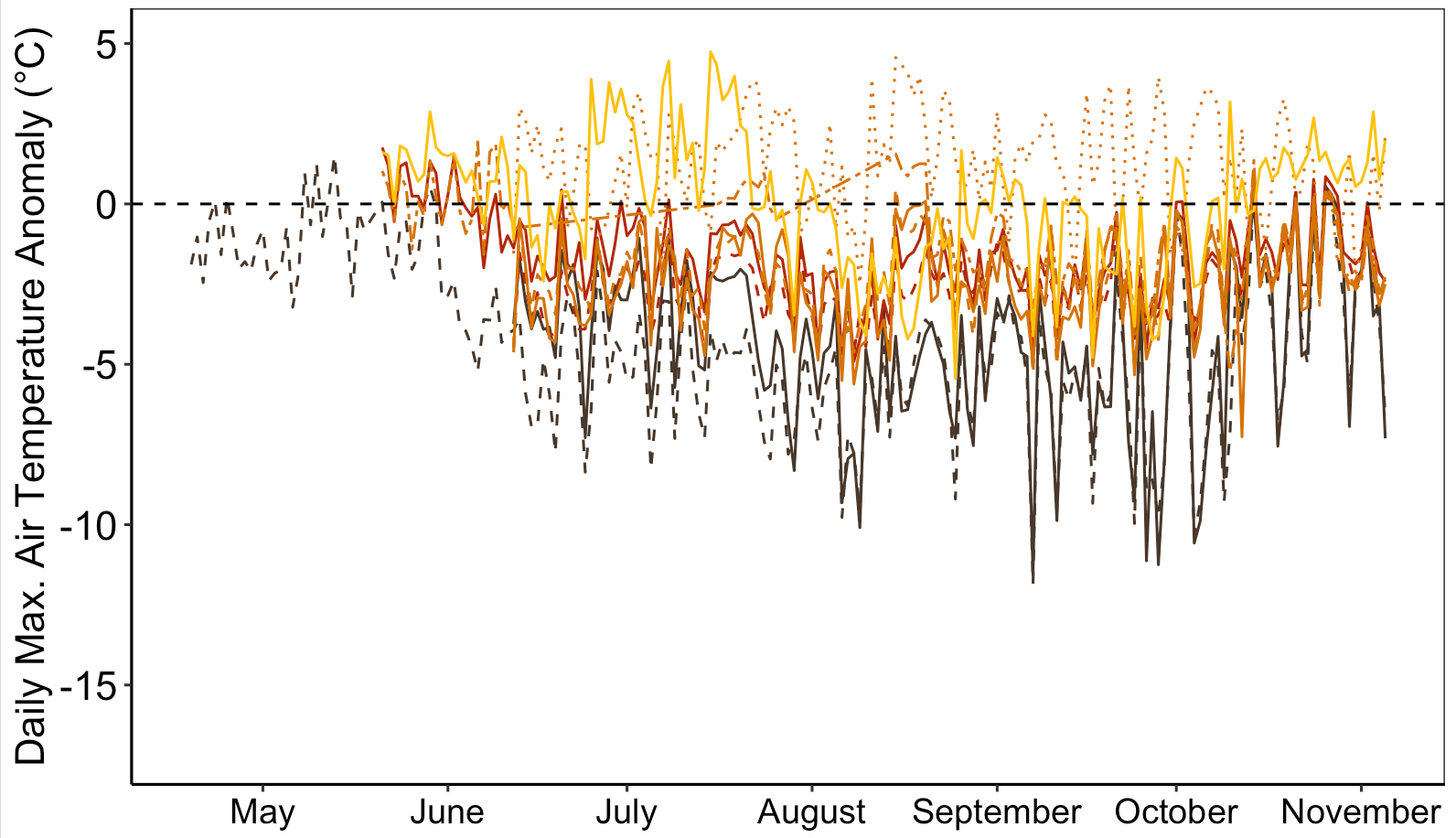

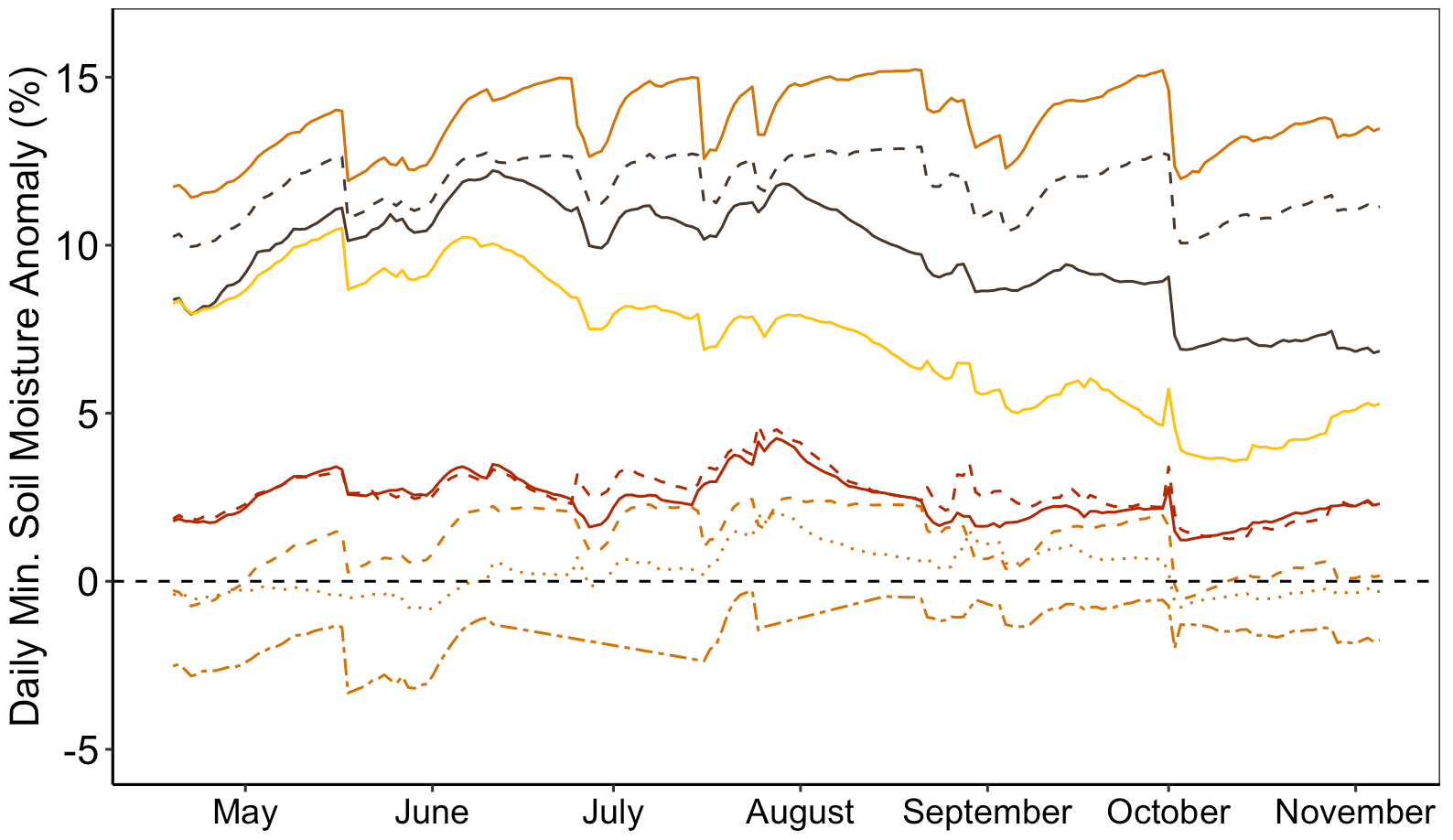

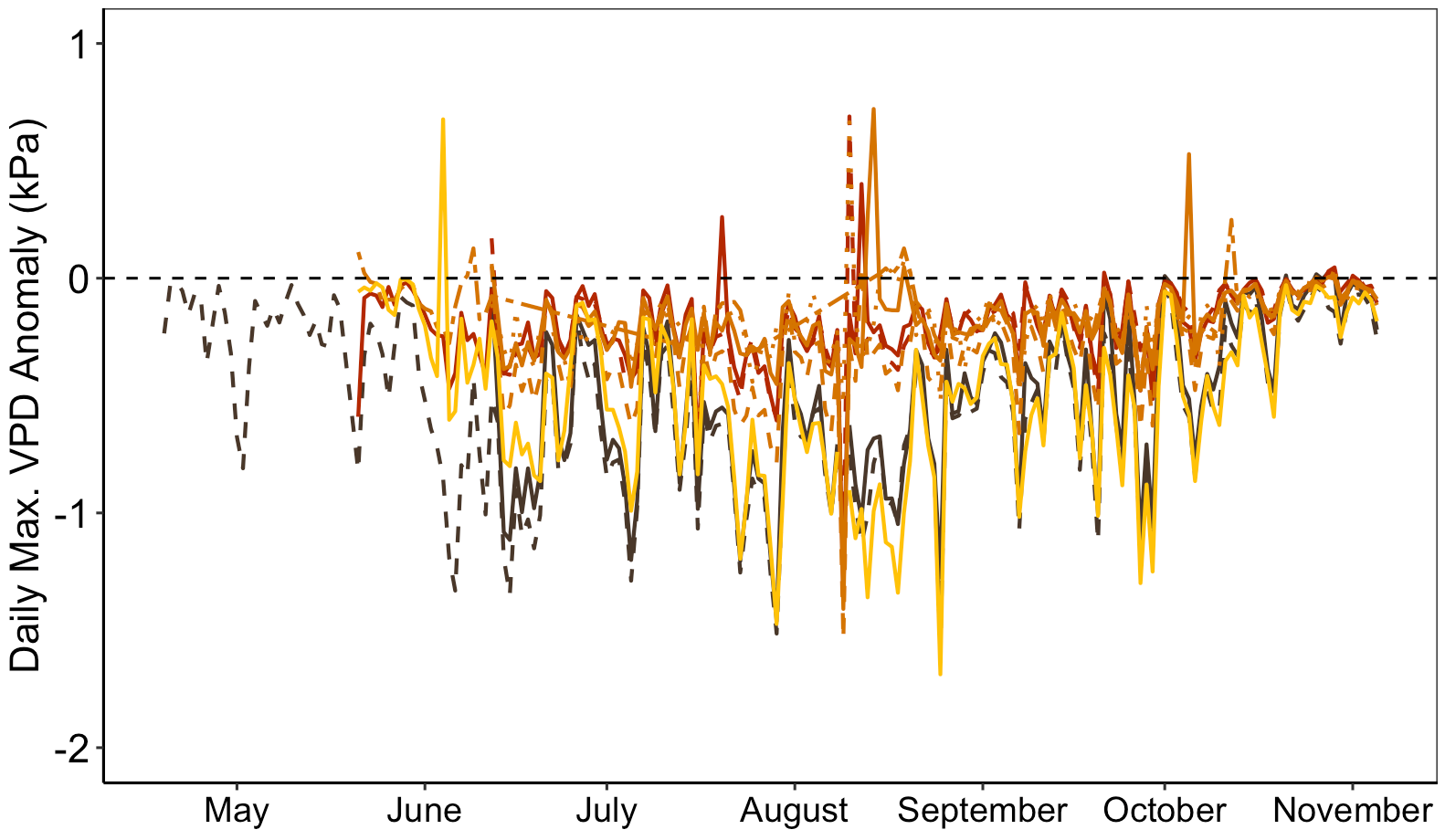

**(d) 2022**

**(a) 2021**

**(b) 2021**

**(c) 2021**

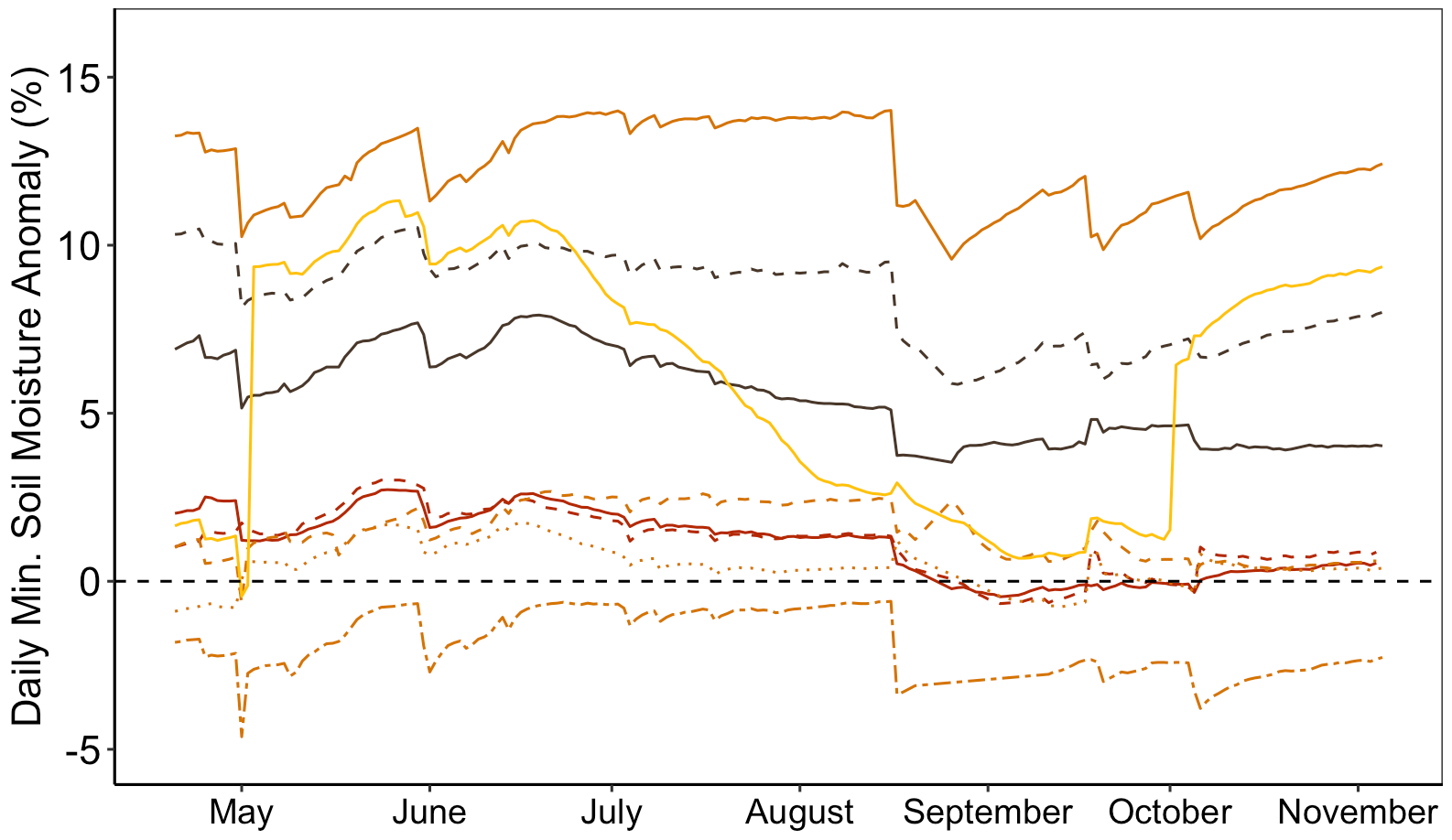

**(e) 2022**

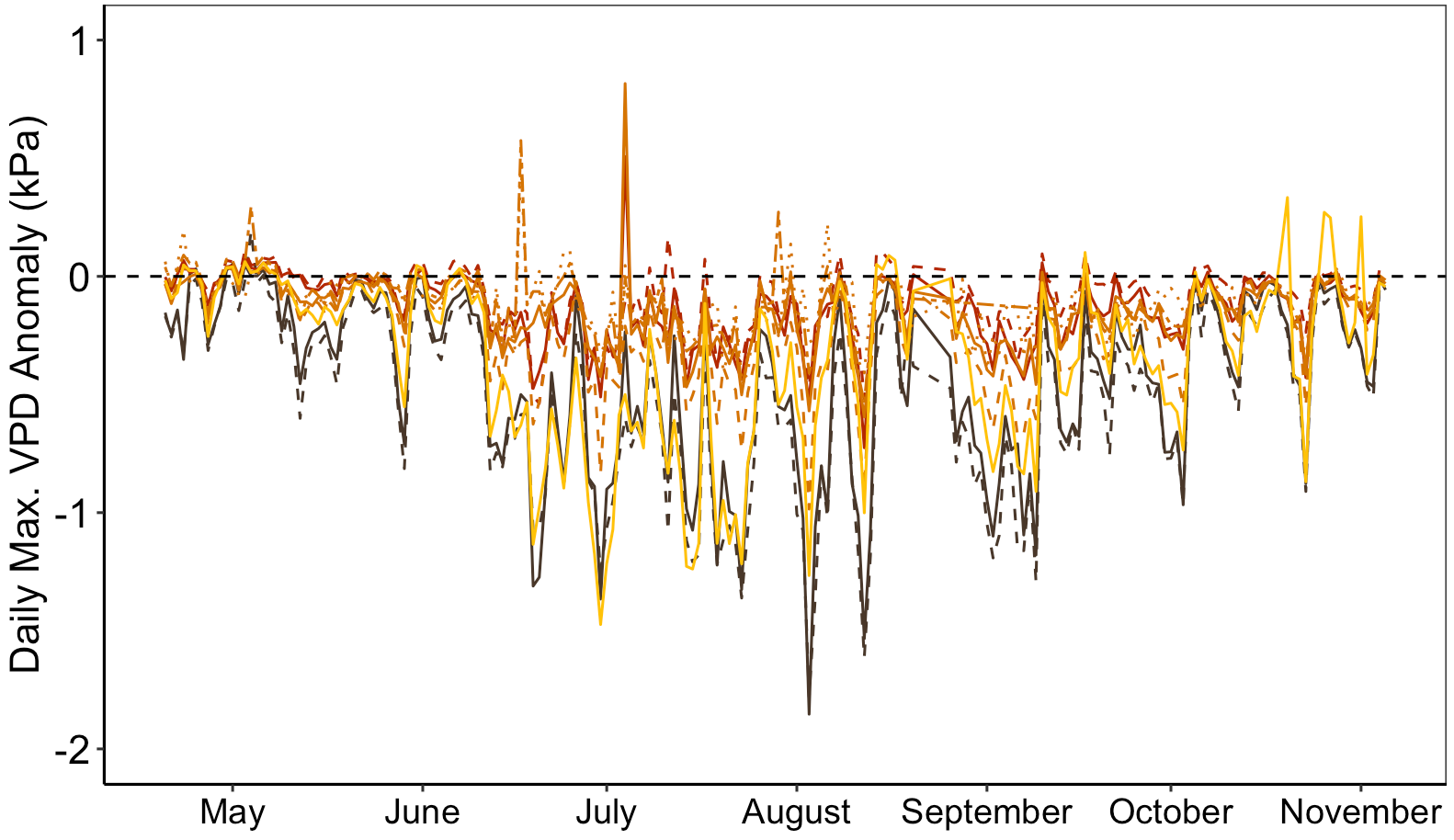

**(f) 2022**

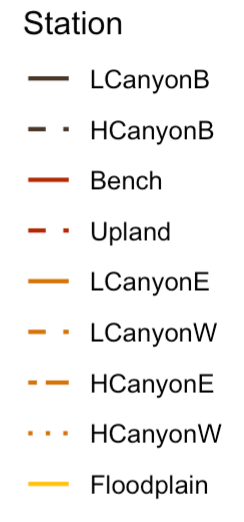

*Buffering*

*Buffering*

*Buffering*

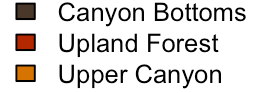

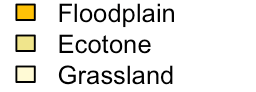

Canyon Bottoms

Upland Forest

Upper Canyon

Floodplain

**Figure S1. Daily microclimate anomalies at nine forested monitoring stations relative to ambient macroclimate conditions in the Niobrara Plot, reproduced from McNichol et al. (2024).** Time series showing the daily (a,d) maximum air temperature anomaly (°C), (b,e) maximum vapor pressure deficit anomaly (kPa), and (c,f) minimum soil moisture anomaly (%) during the growing season (April-November) in 2021 (a-c) and 2022 (d-f) from McNichol et al. (2024), illustrating the degree of microclimate buffering across habitats. Anomalies were calculated by subtracting daily values at an open prairie station representing the regional macroclimate from corresponding values at nine forested stations spanning canyon bottoms, upland forest, upper canyon, and floodplain habitats. Negative anomalies in maximum air temperature and VPD and positive anomalies in minimum soil moisture indicate microclimate buffering; the direction of buffering is labeled in panels (d-f). Lines represent individual monitoring stations color-coded by habitat type; line types distinguish stations within habitats as shown in the inset legend. Canyon bottoms, upland forest, and upper canyon consistently showed the largest buffering anomalies across both years and all three variables and were designated as microrefugial habitats; the floodplain showed partial buffering in soil moisture and VPD but positive temperature anomalies, reflecting its high-light, more exposed conditions.

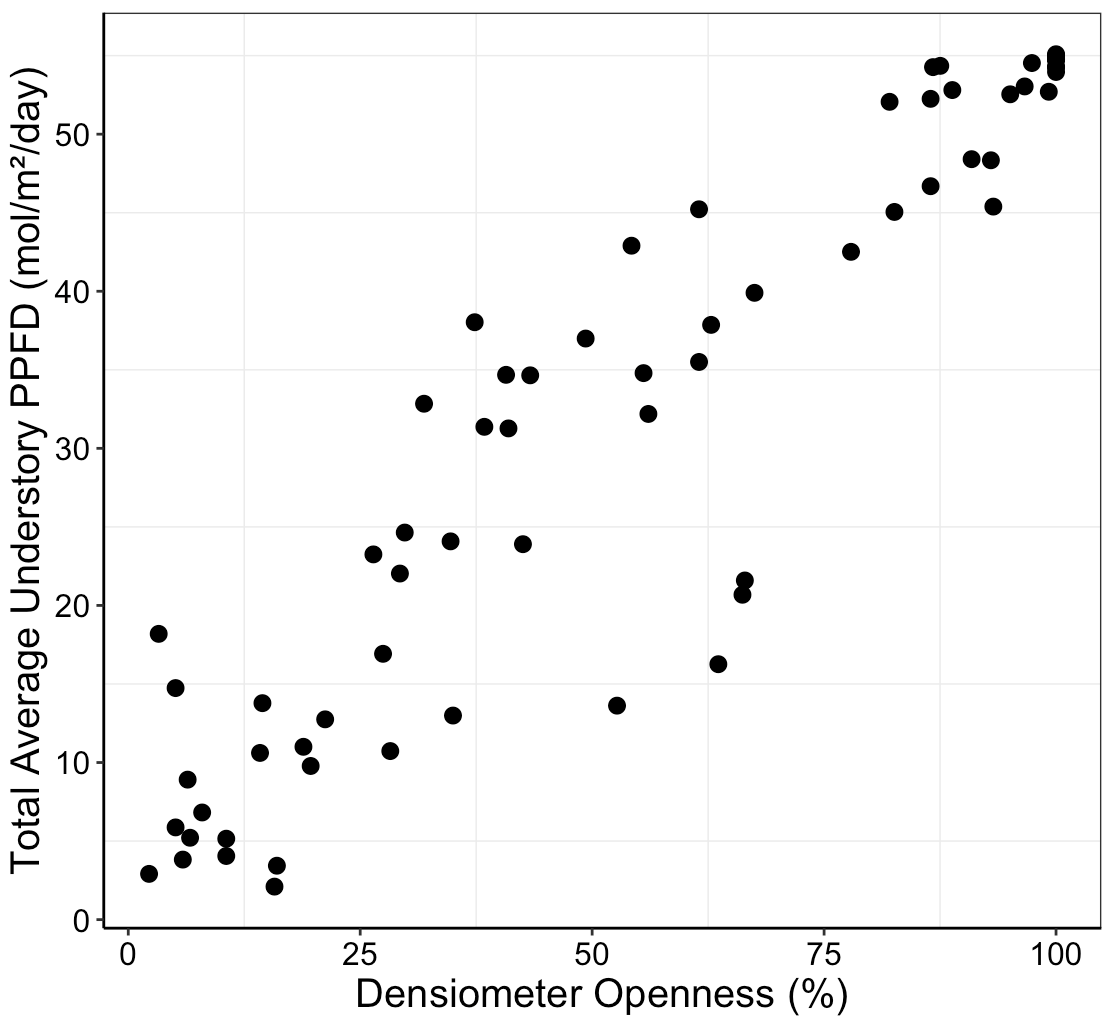

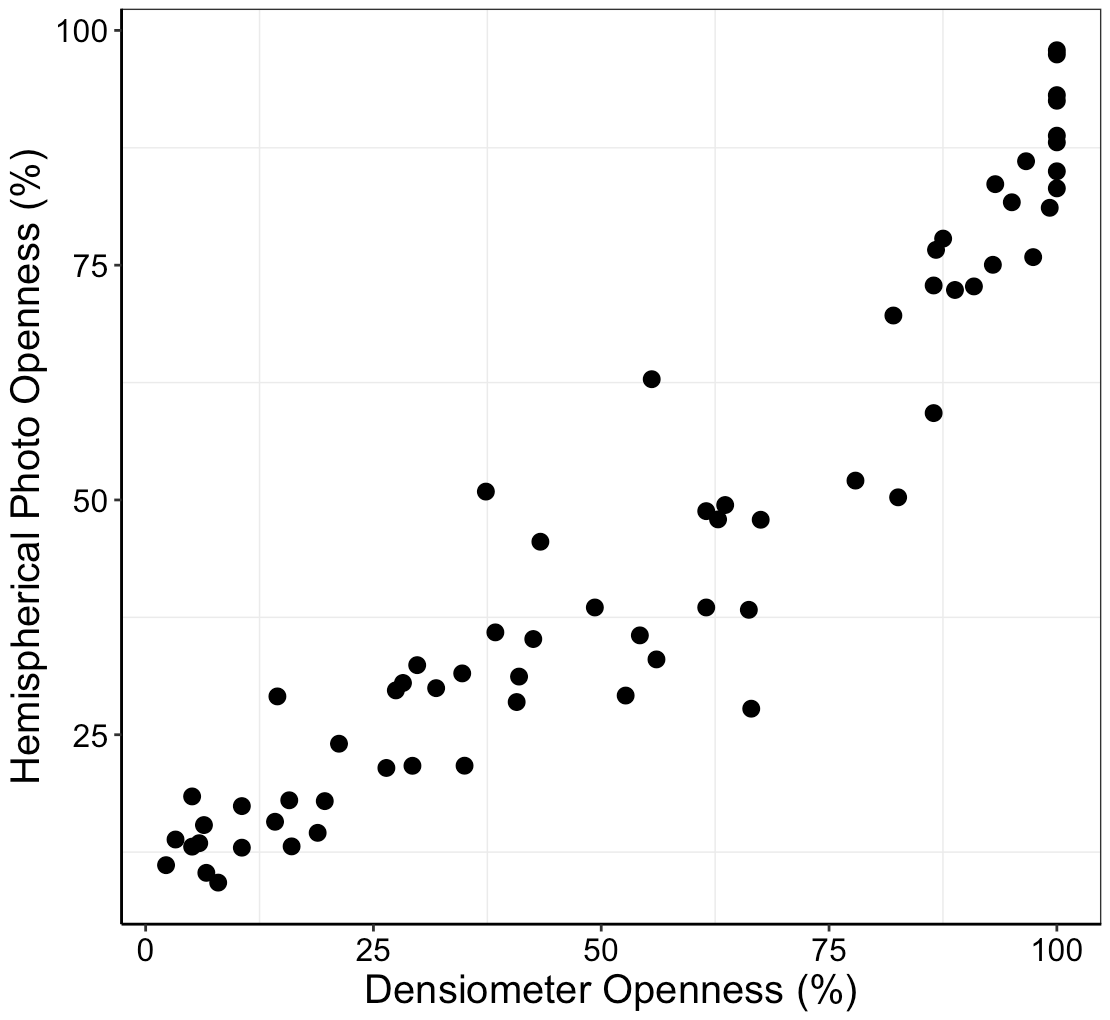

**(a) *r* = 0.95**

**(b) *r* = 0.93**

**Figure S2. Relationship between two estimates of canopy openness, and between canopy openness and the estimated total average daily photosynthetically active photon flux density at 68 subplots in the Niobrara plot.** Scatterplots showing strong positive correlations between densiometer-estimated canopy openness versus (a) canopy openness estimated from hemispherical photographs of the canopy, and (b) total average daily photosynthetically active photon flux density estimated from hemispherical photographs of the canopy. Significant relationships at 𝛼 = 0.05 shown in bold.

**Table S1. Summary of understory woody and herbaceous plant species richness by functional group and with respect to topographic habitats in the Niobrara Plot in 2021**. Numbers represent the total species richness per functional group sampled within each habitat (columns), with the right-most column indicating the total sampled richness per functional group. Numbers for understory woody plants and tree seedlings only reflect richness in the 78 subplots sampled for herbaceous plants in 2021 for direct comparison (see Table S2 for understory woody species richness across all subplots in 2021 and 2022). Since only one subplot sampled for herbaceous plants was in grassland, we grouped this subplot with the ecotone for summary and analyses. The following number of herbaceous subplots were sampled in each habitat type: canyon bottoms = 20; upland forest = 22; upper canyon = 18; floodplain = 9; ecotone = 9.

| **Functional Group** | **Canyon Bottoms** | **Upland Forest** | **Upper Canyon** | **Floodplain** | **Ecotone** | **Total Richness** |
| --- | --- | --- | --- | --- | --- | --- |
| Non-vascular and lower vascular plants | 3 | 0 | 1 | 0 | 0 | 3 |
| Tree seedlings | 10 | 9 | 8 | 8 | 2 | 15 |
| Sedges | 4 | 4 | 5 | 1 | 2 | 7 |
| Forbs | 39 | 51 | 47 | 46 | 24 | 90 |
| Understory woody plants | 6 | 4 | 5 | 2 | 0 | 7 |
| C_3_ grasses | 5 | 9 | 6 | 10 | 10 | 17 |
| C_4_ grasses | 2 | 8 | 6 | 10 | 17 | 22 |

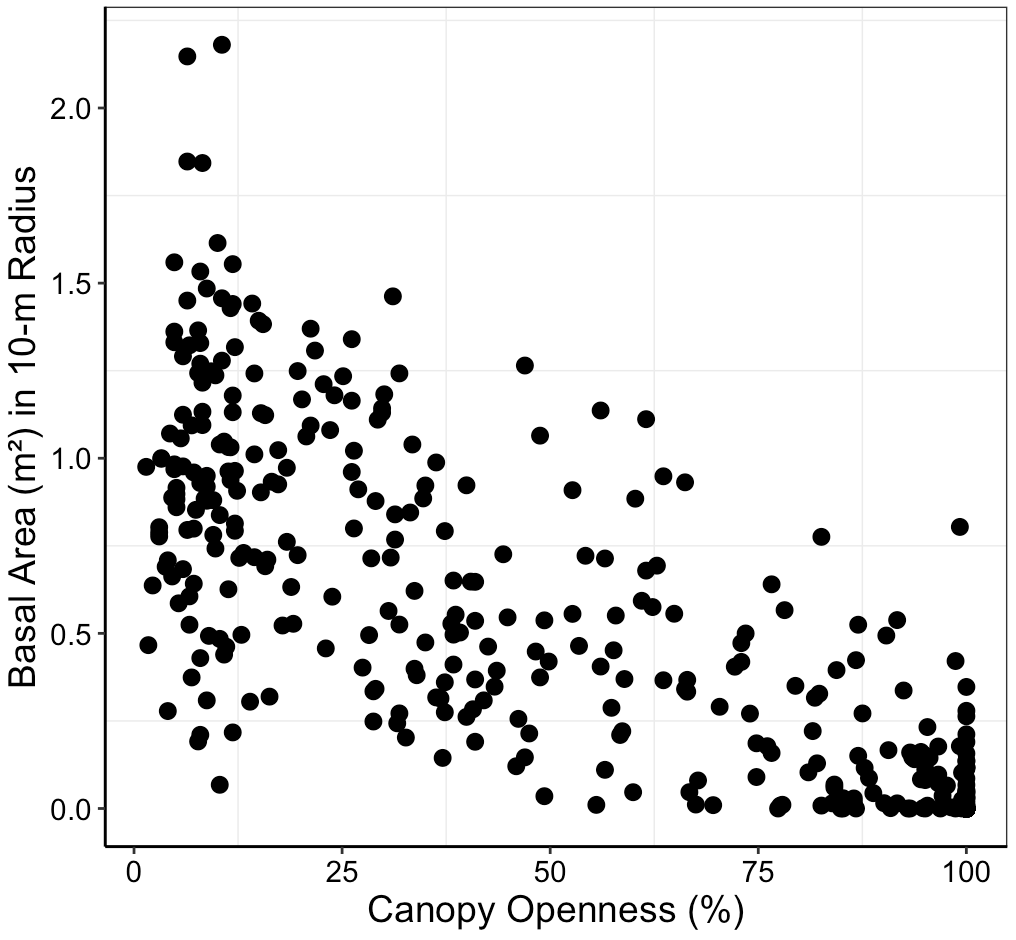

***r* = -0.83**

**Figure S3. Relationship between canopy openness and overstory (DBH ≥1 cm) basal area around each subplot in the Niobrara plot.** Scatterplot showing statistically significant ($\alpha$ = 0.05) negative correlation between densiometer-estimated canopy openness and the total overstory basal area in the 10-m radius around each of the 504 1 × 1 m subplots.

**Table S2. Taxonomy, growth form, occurrence, and abundance of 31 understory woody species documented in 2021 and 2022 in the 20.2 ha Niobrara Forest Dynamics Plot, North-Central Nebraska, USA.** Understory woody plants were sampled in 504 1 × 1 m subplots within the Niobrara Plot (Figure 1). Occurrence is the total number of subplots in which a species was documented, and abundance is the total number of censused stems summed across all subplots. *Observations of *Parthenocissus quinquefolia* and *P. vitacea* were grouped due to difficulty distinguishing them and were recorded as present or absent due to the difficulty in distinguishing genets from ramets aboveground.

| **Species (Family)** | **Growth form** | **2021 occurrence** | **2021 abundance** | **2022 occurrence** | **2022 abundance** |
| --- | --- | --- | --- | --- | --- |
| *Amorpha nana* Nutt. (Fabaceae) | Shrub | 0 | 0 | 1 | 2 |
| *Betula papyrifera* Marshall (Betulaceae) | Canopy tree | 1 | 1 | 0 | 0 |
| *Celastrus scandens* L. (Celastraceae) | Liana | 16 | 43 | 16 | 53 |
| *Celtis occidentalis* L. (Cannabaceae) | Canopy tree | 116 | 391 | 117 | 424 |
| *Fraxinus pennsylvanica* Marshall (Oleaceae) | Canopy tree | 56 | 144 | 58 | 143 |
| *Juglans nigra* L. (Juglandaceae) | Canopy tree | 6 | 6 | 8 | 8 |
| *Juniperus virginiana* L. (Cupressaceae) | Canopy tree | 45 | 111 | 49 | 110 |
| *Lonicera dioica* L. (Caprifoliaceae) | Liana | 2 | 14 | 2 | 13 |
| *Ostrya virginiana* (Mill.) K. Koch (Betulaceae) | Subcanopy tree | 69 | 225 | 67 | 251 |
| *Parthenocissus quinquefolia* L. Planch. and *P. vitacea* (Knerr) Hitchcock (Vitaceae) | Liana | 64 | NA* | 68 | NA* |
| *Physocarpus opulifolius* (L.) Maxim. (Rosaceae) | Shrub | 2 | 6 | 1 | 2 |
| *Pinus ponderosa* Lawson & C. Lawson (Pinaceae) | Canopy tree | 1 | 2 | 1 | 2 |
| *Populus deltoides* W. Bartram ex Marshall (Salicaceae) | Canopy tree | 1 | 1 | 2 | 2 |
| *Prunus americana* Marshall (Rosaceae) | Understory tree | 6 | 28 | 8 | 24 |
| *Prunus virginiana* L. (Rosaceae) | Understory tree | 12 | 40 | 11 | 45 |
| *Quercus macrocarpa* Michx. (Fagaceae) | Canopy tree | 31 | 40 | 27 | 33 |
| *Rhus aromatica* Aiton (Anacardiaceae) | Shrub | 2 | 5 | 2 | 6 |
| *Rhus glabra* L. (Anacardiaceae) | Understory tree | 131 | 488 | 124 | 421 |
| *Ribes americanum* Mill. (Grossulariaceae) | Shrub | 1 | 1 | 1 | 1 |
| *Rosa arkansana* Porter (Rosaceae) | Shrub | 13 | 62 | 15 | 66 |
| *Rubus occidentalis* L. (Rosaceae) | Shrub | 87 | 363 | 84 | 360 |
| *Salix amygdaloides* Andersson (Salicaceae) | Understory tree | 2 | 6 | 2 | 5 |
| *Salix eriocephala* Michx. (Salicaceae) | Understory tree | 1 | 5 | 1 | 3 |
| *Salix interior* Rowlee (Salicaceae) | Understory tree | 4 | 26 | 4 | 29 |
| *Symphoricarpos occidentalis* (R.Br.) Hook. (Caprifoliaceae) | Shrub | 20 | 133 | 21 | 150 |
| *Tilia americana* L. (Malvaceae) | Canopy tree | 88 | 419 | 83 | 511 |
| *Toxicodendron rydbergii* (Small) Greene (Anacardiaceae) | Shrub | 29 | 189 | 27 | 189 |
| *Ulmus rubra* Muhl. (Ulmaceae) | Canopy tree | 10 | 18 | 8 | 18 |
| *Vitis riparia* Michx. (Vitaceae) | Liana | 71 | 200 | 58 | 129 |
| *Zanthoxylum americanum* Mill. (Rutaceae) | Shrub | 4 | 4 | 4 | 5 |

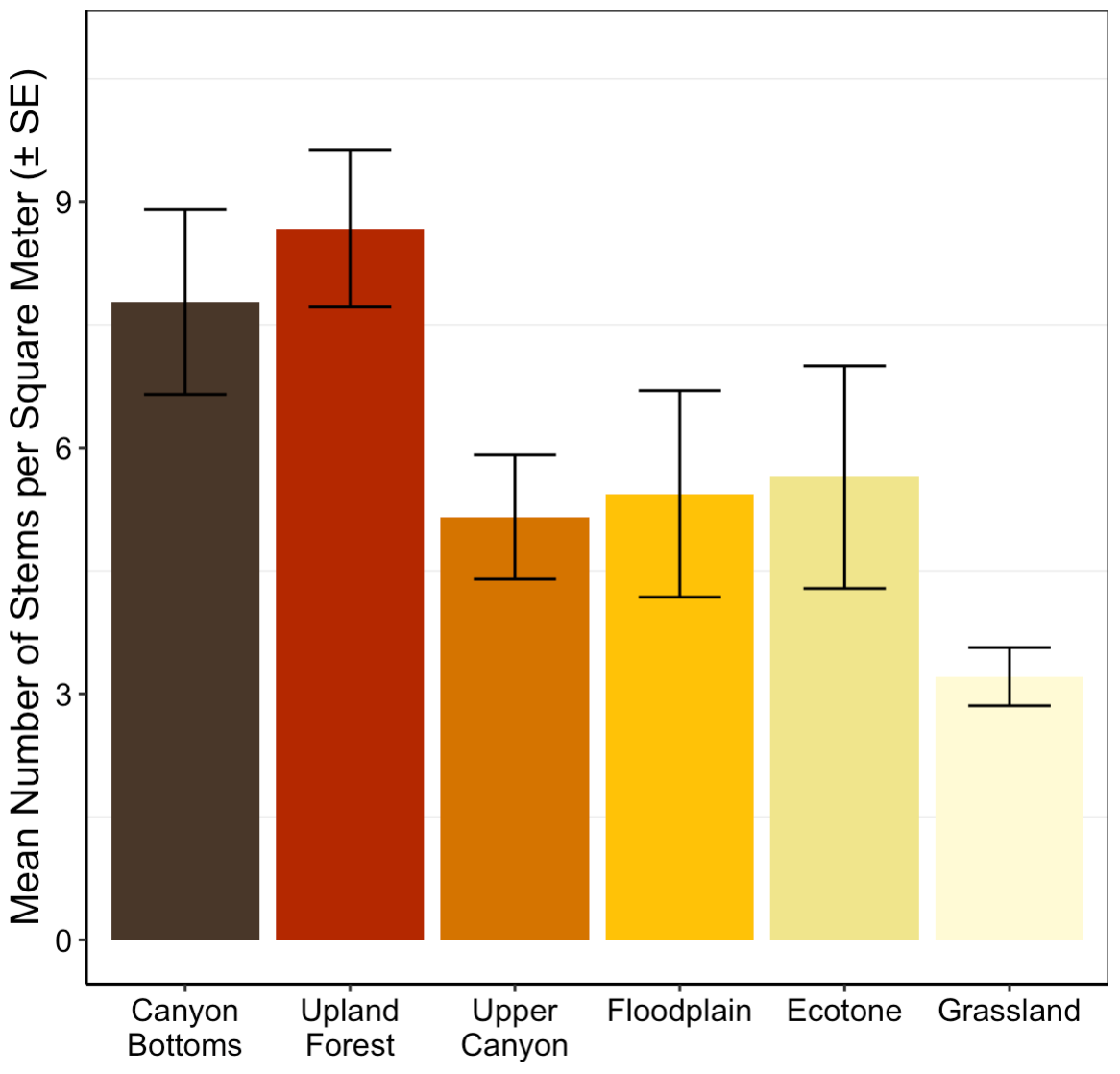

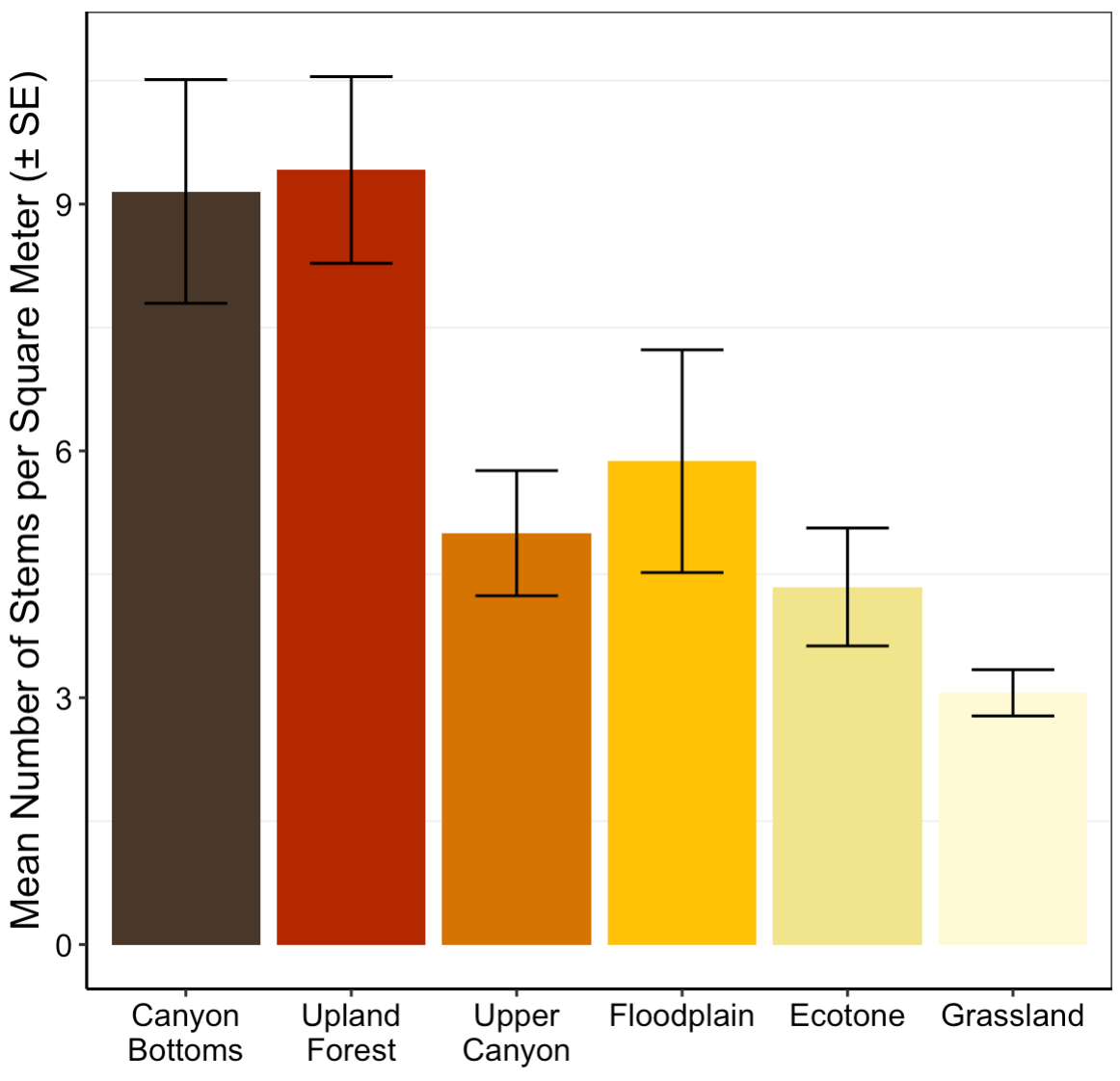

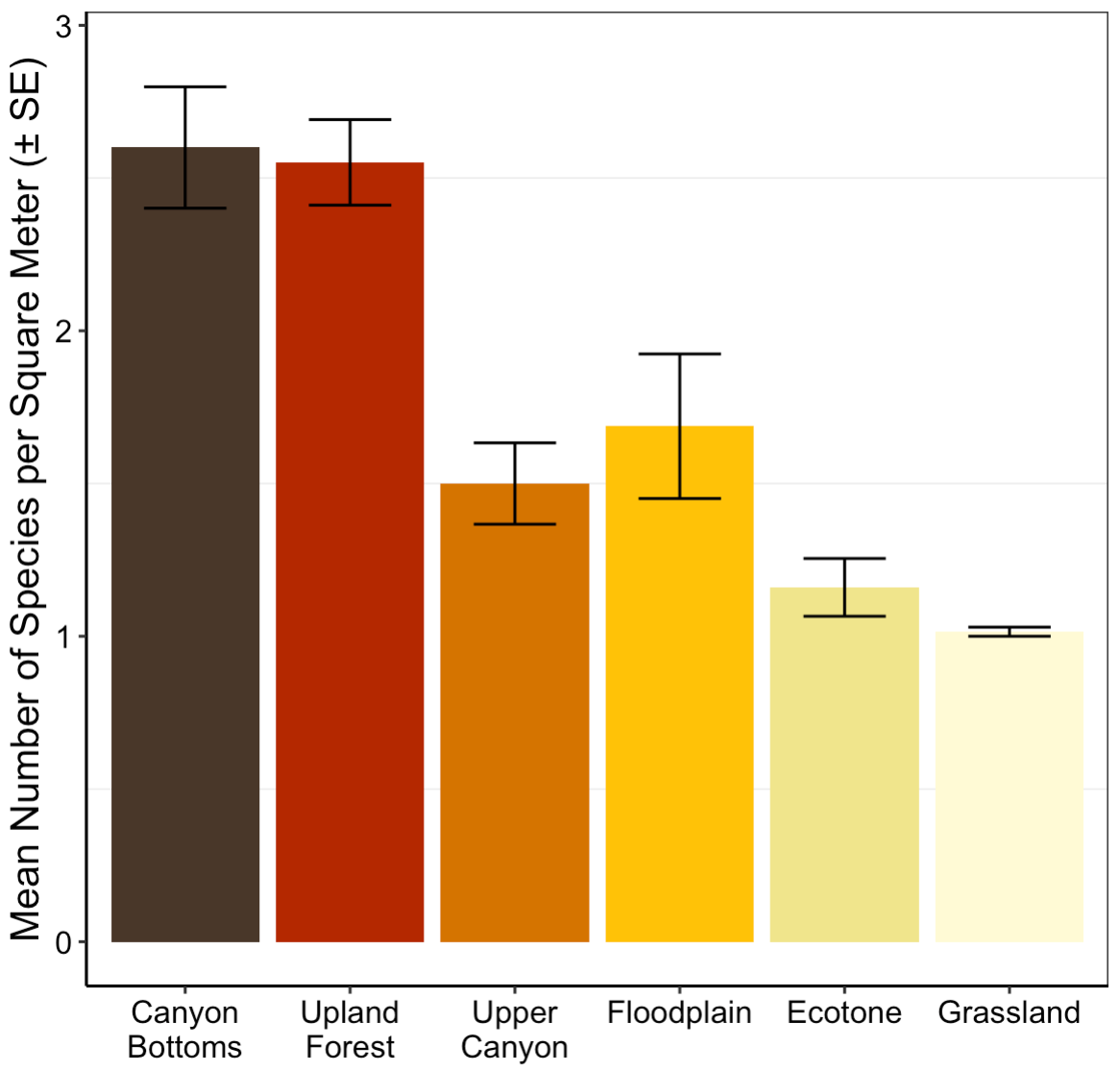

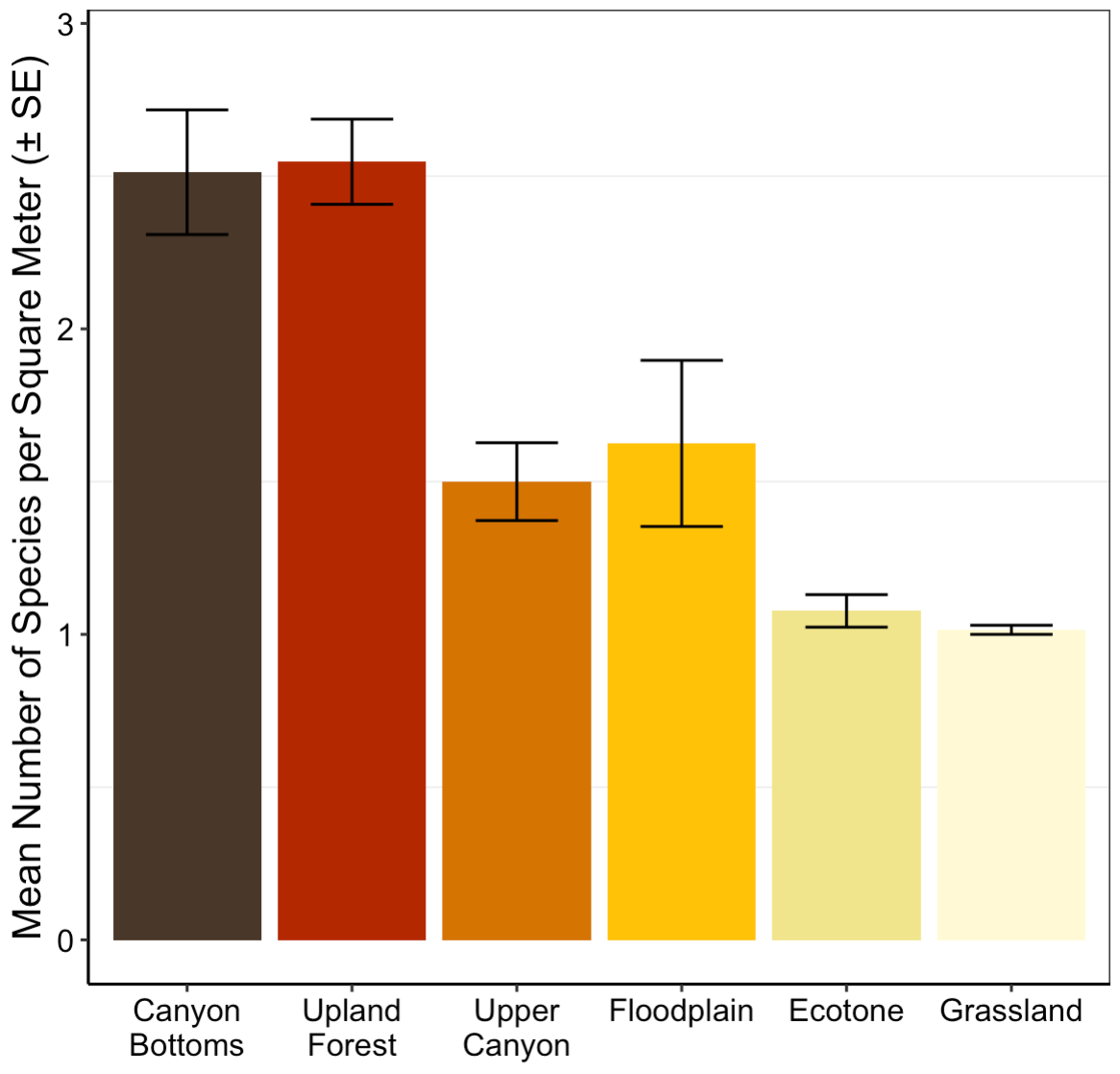

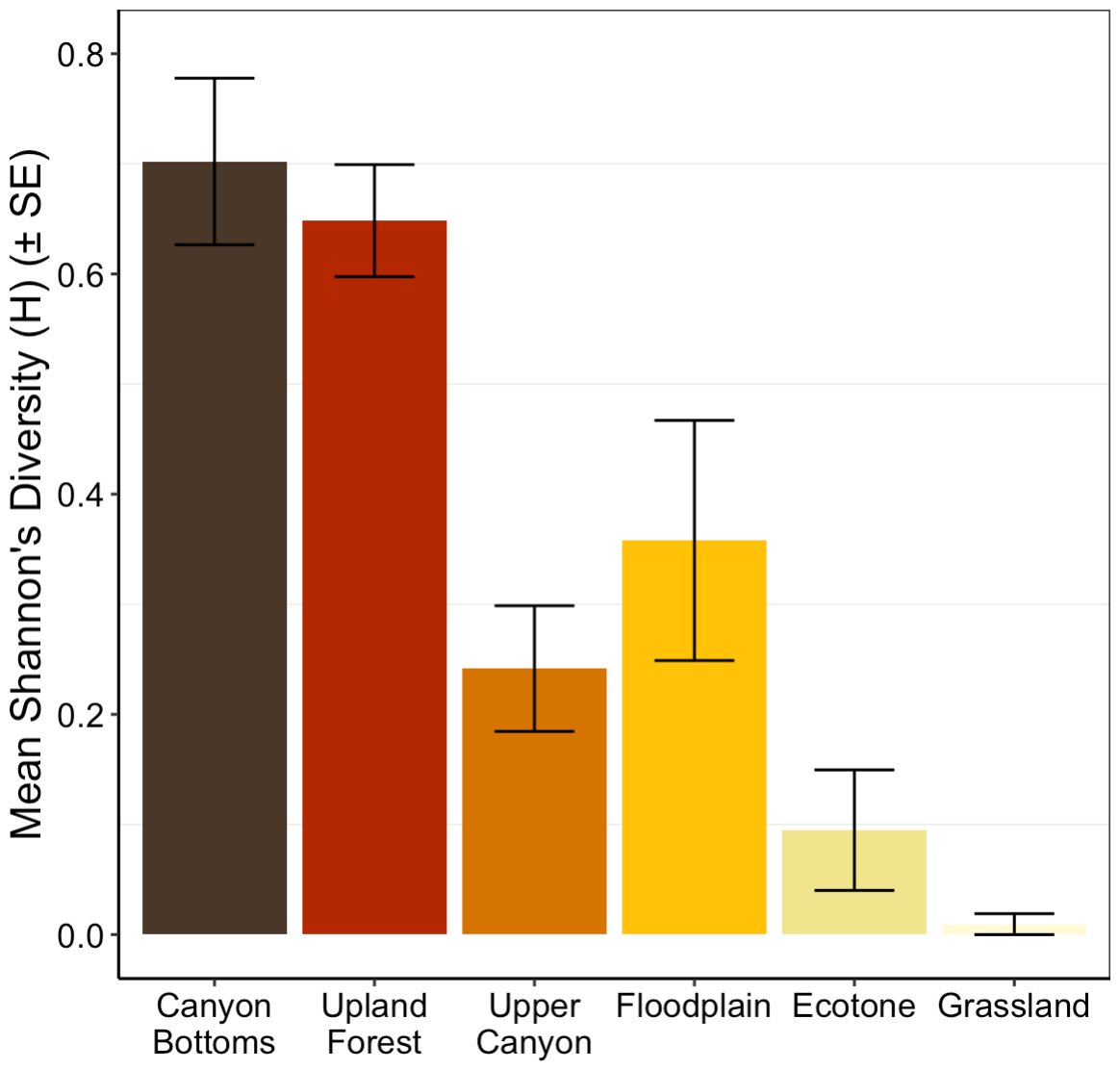

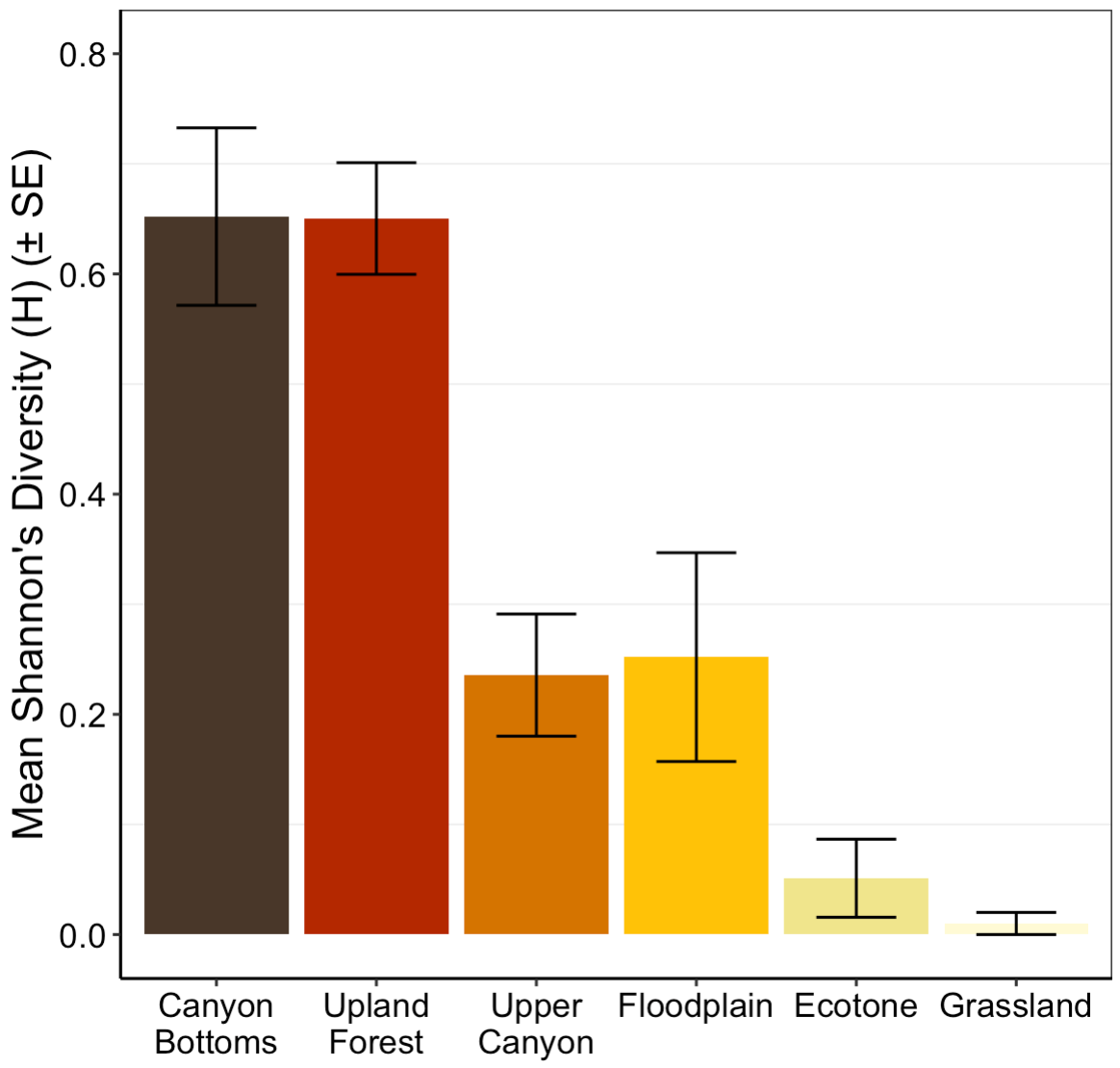

**ab**

**a**

**bc**

**abc**

**abc**

**c**

**a**

**a**

**ab**

**ab**

**bd**

**cd**

**a**

**b**

**c**

**a**

**bc**

**b**

**a**

**a**

**a**

**ab**

**b**

**c**

**a**

**bc**

**a**

**b**

**b**

**c**

**a**

**bc**

**(c) 2021**

**(d) 2022**

**(a) 2021**

**(b) 2022**

**(f) 2022**

**(e) 2021**

**b**

**b**

**c**

**bc**

**Figure S4. Variation in tree seedling stem density, species richness, and Shannon’s diversity among six habitats in the Niobrara plot.** Barplots show habitat-related variation in mean (± standard error, SE) tree seedling (a,b) stem density, (c,d) species richness, and (e,f) Shannon’s Diversity (H) per square meter in 2021 (a,c,e) and 2022 (b,d,f). Habitats are ordered by increasing solar radiation and exposure from canyon bottoms to grassland. Letters above the bars indicate significant pairwise differences at $\alpha$ = 0.05 after correction for multiple comparisons using Tukey’s Honestly Significant Differences.

**Table S3. Summary statistics from one-way analyses of variance testing for differences among habitats in understory woody plant and tree seedling stem density, species richness, and Shannon’s diversity index in the Niobrara Plot in 2021 and 2022**. Models were fit separately for all understory woody plants and for tree seedlings only. DF = degrees of freedom; significant effects at $\alpha$ = 0.05 are shown in bold.

| **Understory Woody Plants** | | | | | |
| --- | --- | --- | --- | --- | --- |
| **Year** | **Response Variable** | ***F*-value** | **DF (Error, Total)** | ***R*^2^** | ***p*-value** |
| 2021 | Stem density | 9.805 | 5, 350 | 0.110 | **<0.001** |
| 2022 | Stem density | 10.860 | 5, 341 | 0.125 | **<0.001** |
| 2021 | Species richness | 33.500 | 5, 350 | 0.314 | **<0.001** |
| 2022 | Species richness | 31.730 | 5, 341 | 0.308 | **<0.001** |
| 2021 | Shannon’s diversity | 31.260 | 5, 350 | 0.299 | **<0.001** |
| 2022 | Shannon’s diversity | 30.150 | 5, 341 | 0.296 | **<0.001** |
| **Tree Seedlings** | | | | | |
| **Year** | **Response Variable** | ***F*-value** | **DF (Error, Total)** | ***R*^2^** | ***p*-value** |
| 2021 | Stem density | 6.160 | 5, 301 | 0.078 | **<0.001** |
| 2022 | Stem density | 7.383 | 5, 300 | 0.095 | **<0.001** |
| 2021 | Species richness | 32.720 | 5, 301 | 0.341 | **<0.001** |
| 2022 | Species richness | 30.990 | 5, 300 | 0.330 | **<0.001** |
| 2021 | Shannon’s diversity | 27.880 | 5, 301 | 0.305 | **<0.001** |
| 2022 | Shannon’s diversity | 28.800 | 5, 300 | 0.313 | **<0.001** |

**Table S4. Summary statistics from one-way analyses of variance evaluating differences among habitats in proportional species richness of herbaceous plant functional groups and the ratio of woody to non-woody species (rWNW) in the Niobrara Plot in 2021.** Non-vascular and lower vascular plants were not statistically analyzed due to few observations across subplots. DF = degrees of freedom; significant effects at $\alpha$ = 0.05 shown in bold.

| **Functional group** | **Predictor** | ***F*-value** | **DF (Error, Total)** | ***R*^2^** | ***p*-value** |
| --- | --- | --- | --- | --- | --- |
| Sedges | habitat type | 2.451 | 4, 72 | 0.120 | **0.054** |
| C_3_ grasses | habitat type | 1.543 | 4, 72 | 0.079 | 0.199 |
| C_4_ grasses | habitat type | 19.480 | 4, 72 | 0.520 | **<0.001** |
| Forbs | habitat type | 0.217 | 4, 72 | 0.012 | 0.928 |
| rWNW | habitat type | 3.253 | 4, 71 | 0.155 | **0.016** |

**Table S5. Summary statistics for generalized additive models (GAMs) evaluating effects of continuous topography and canopy openness on understory woody plant and tree seedling stem density and species richness in the Niobrara Plot in 2021 and 2022.** Prior to model fitting, a principal components analysis (PCA) was conducted on elevation, slope, northness, and eastness, and the first two topographic PC axes (topoPC1, topoPC2) were retained as topographic predictors. GAMs were fit with topoPC1, topoPC2, and openness as predictors, with a two-dimensional thin-plate spline of subplot coordinates included to account for spatial autocorrelation. Adjusted *R*^2^ values, parameter estimates, standard errors (SE), *t*-values, and *p*-values are reported; significant effects at $\alpha$ = 0.05 are shown in bold. Parameters have not been back transformed.

|  | **Understory Woody Plants** | | | | | | |
| --- | --- | --- | --- | --- | --- | --- | --- |
|  | **Year** | **Response Variable** | **Model Term** | **Estimate** | **SE** | ***t*-value** | ***p*-value** |
|  | 2021 | Stem density | Intercept | 1.862 | 0.135 | 13.733 | **<0.001** |
|  |  | *R*^2^ = 0.30 | topoPC1 | 0.141 | 0.077 | 1.828 | 0.068 |
|  |  |  | topoPC2 | -0.071 | 0.054 | -1.295 | 0.196 |
|  |  |  | openness | -0.005 | 0.002 | -2.489 | **0.013** |
|  | 2022 | Stem density | Intercept | 1.916 | 0.139 | 13.731 | **<0.001** |
|  |  | *R*^2^ = 0.29 | topoPC1 | 0.114 | 0.079 | 1.434 | 0.153 |
|  |  |  | topoPC2 | -0.057 | 0.056 | -1.012 | 0.312 |
|  |  |  | openness | -0.006 | 0.002 | -2.618 | **0.009** |
|  | 2021 | Species richness | Intercept | 0.647 | 0.025 | 26.173 | **<0.001** |
|  |  | *R*^2^ = 0.49 | topoPC1 | -0.021 | 0.014 | -1.511 | 0.132 |
|  |  |  | topoPC2 | -0.004 | 0.010 | -0.354 | 0.723 |
|  |  |  | openness | 0.002 | <0.001 | 6.184 | **<0.001** |
|  | 2022 | Species richness | Intercept | 0.666 | 0.025 | 26.761 | **<0.001** |
|  |  | *R*^2^ = 0.49 | topoPC1 | -0.031 | 0.014 | -2.148 | **0.032** |
|  |  |  | topoPC2 | -0.006 | 0.010 | -0.554 | 0.580 |
|  |  |  | openness | 0.002 | <0.001 | 5.299 | **<0.001** |
|  | **Tree Seedlings** | | | | | | |
|  | **Year** | **Response Variable** | **Model Term** | **Estimate** | **SE** | ***t*-value** | ***p*-value** |
| 2021 | | Stem density | Intercept | 1.710 | 0.139 | 12.292 | **<0.001** |
|  | | *R*^2^ = 0.29 | topoPC1 | -0.020 | 0.082 | -0.244 | 0.808 |
|  | |  | topoPC2 | 0.030 | 0.058 | 0.520 | 0.603 |
|  | |  | openness | -0.007 | 0.002 | -3.146 | **0.002** |
| 2022 | | Stem density | Intercept | 1.828 | 0.143 | 12.758 | **<0.001** |
|  | | *R*^2^ = 0.31 | topoPC1 | -0.042 | 0.082 | -0.511 | 0.610 |
|  | |  | topoPC2 | 0.023 | 0.057 | 0.410 | 0.682 |
|  | |  | openness | -0.009 | 0.002 | -3.861 | **<0.001** |
| 2021 | | Species richness | Intercept | 0.554 | 0.035 | 15.642 | **<0.001** |
|  | | *R*^2^ = 0.54 | topoPC1 | -0.027 | 0.021 | -1.314 | 0.190 |
|  | |  | topoPC2 | -0.021 | 0.015 | -1.400 | 0.163 |
|  | |  | openness | 0.003 | <0.001 | 6.023 | **<0.001** |
| 2022 | | Species richness | Intercept | 0.535 | 0.044 | 12.065 | **<0.001** |
|  | | *R*^2^ = 0.54 | topoPC1 | -0.055 | 0.025 | -2.195 | **0.029** |
|  | |  | topoPC2 | -0.020 | 0.018 | -1.130 | 0.260 |
|  | |  | openness | 0.003 | <0.001 | 4.269 | **<0.001** |

**Table S6. Summary statistics for zero-inflated negative binomial models evaluating the effects of continuous topography and canopy openness on the proportional species richness of herbaceous plant functional groups and the ratio of woody to non-woody species (rWNW) in the Niobrara Plot in 2021.** Prior to model fitting, a principal components analysis (PCA) was conducted on elevation, slope, northness, and eastness; the first two PC axes (topoPC1, topoPC2) were retained as topographic predictors alongside openness. Parameter estimates, standard errors (SE), *z*-values, and *p*-values are shown separately for the count component (species richness conditional on presence) and the zero-inflation component (probability of absence). McFadden's pseudo-*R*² (p*R*²) is reported for each model; significant effects at α = 0.05 are shown in bold. *Forbs were modeled using a binomial generalized linear model with a single model component; therefore, presence/absence columns are not applicable (—).

|  |  |  | **Count component (richness \| present)** | | | | **Presence/absence component** | | | |
| --- | --- | --- | --- | --- | --- | --- | --- | --- | --- | --- |
| **Functional group** | ***pR*^2^** | **Predictor** | **Estimate** | **SE** | ***z*-value** | ***p*-value** | **Estimate** | **SE** | ***z*-value** | ***p*-value** |
| Sedges | 0.053 | Intercept | 2.441 | 0.133 | 18.276 | **<0.001** | -3.592 | 0.937 | -3.835 | **<0.001** |
|  |  | topoPC1 | -0.024 | 0.081 | -0.301 | 0.763 | 1.126 | 0.491 | 2.293 | **0.022** |
|  |  | topoPC2 | -0.066 | 0.077 | -0.860 | 0.390 | 0.362 | 0.302 | 1.197 | 0.231 |
|  |  | openness | -0.004 | 0.003 | -1.606 | 0.108 | 0.043 | 0.013 | 3.448 | **<0.001** |
| C_3_ Grasses | 0.021 | Intercept | 2.314 | 0.182 | 12.732 | **<0.001** | -1.114 | 0.608 | -1.831 | 0.067 |
|  |  | topoPC1 | -0.106 | 0.114 | -0.931 | 0.352 | 0.066 | 0.374 | 0.175 | 0.861 |
|  |  | topoPC2 | -0.007 | 0.083 | -0.089 | 0.929 | 0.008 | 0.307 | 0.027 | 0.979 |
|  |  | openness | 0.005 | 0.003 | 2.050 | **0.040** | -0.006 | 0.010 | -0.571 | 0.568 |
| C_4_ Grasses | 0.165 | Intercept | 2.454 | 0.215 | 11.427 | **<0.001** | 1.390 | 0.594 | 2.340 | **0.019** |
|  |  | topoPC1 | -0.403 | 0.101 | -3.977 | **<0.001** | 0.877 | 0.402 | 2.179 | **0.029** |
|  |  | topoPC2 | 0.093 | 0.082 | 1.124 | 0.261 | 0.077 | 0.342 | 0.224 | 0.823 |
|  |  | openness | 0.004 | 0.003 | 1.525 | 0.127 | -0.043 | 0.011 | -3.879 | **<0.001** |
| rWNW | 0.040 | Intercept | 2.937 | 0.179 | 16.420 | **<0.001** | 0.099 | 0.519 | 0.191 | 0.849 |
|  |  | topoPC1 | -0.238 | 0.119 | -2.006 | **0.045** | -0.607 | 0.327 | -1.858 | 0.063 |
|  |  | topoPC2 | -0.023 | 0.089 | -0.257 | 0.797 | 0.032 | 0.265 | 0.120 | 0.905 |
|  |  | openness | -0.005 | 0.003 | -1.643 | 0.100 | 0.008 | 0.008 | 0.928 | 0.353 |
| Forbs* | 0.039 | Intercept | -0.656 | 0.131 | -5.012 | **<0.001** | — | — | — | — |
|  |  | topoPC1 | 0.147 | 0.079 | 1.857 | 0.063 | — | — | — | — |
|  |  | topoPC2 | -0.083 | 0.062 | -1.336 | 0.181 | — | — | — | — |
|  |  | openness | 0.003 | 0.002 | 1.458 | 0.145 | — | — | — | — |

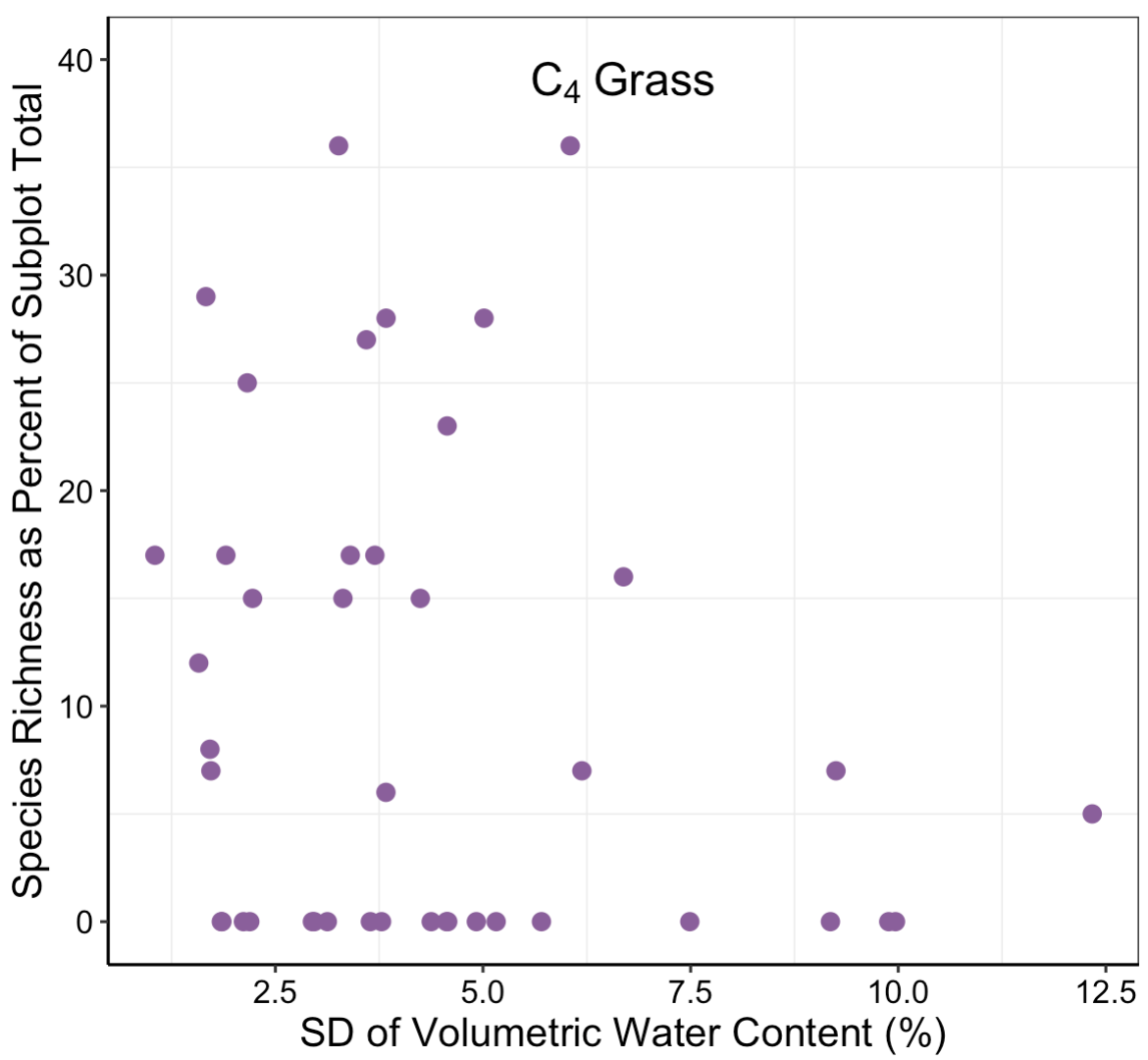

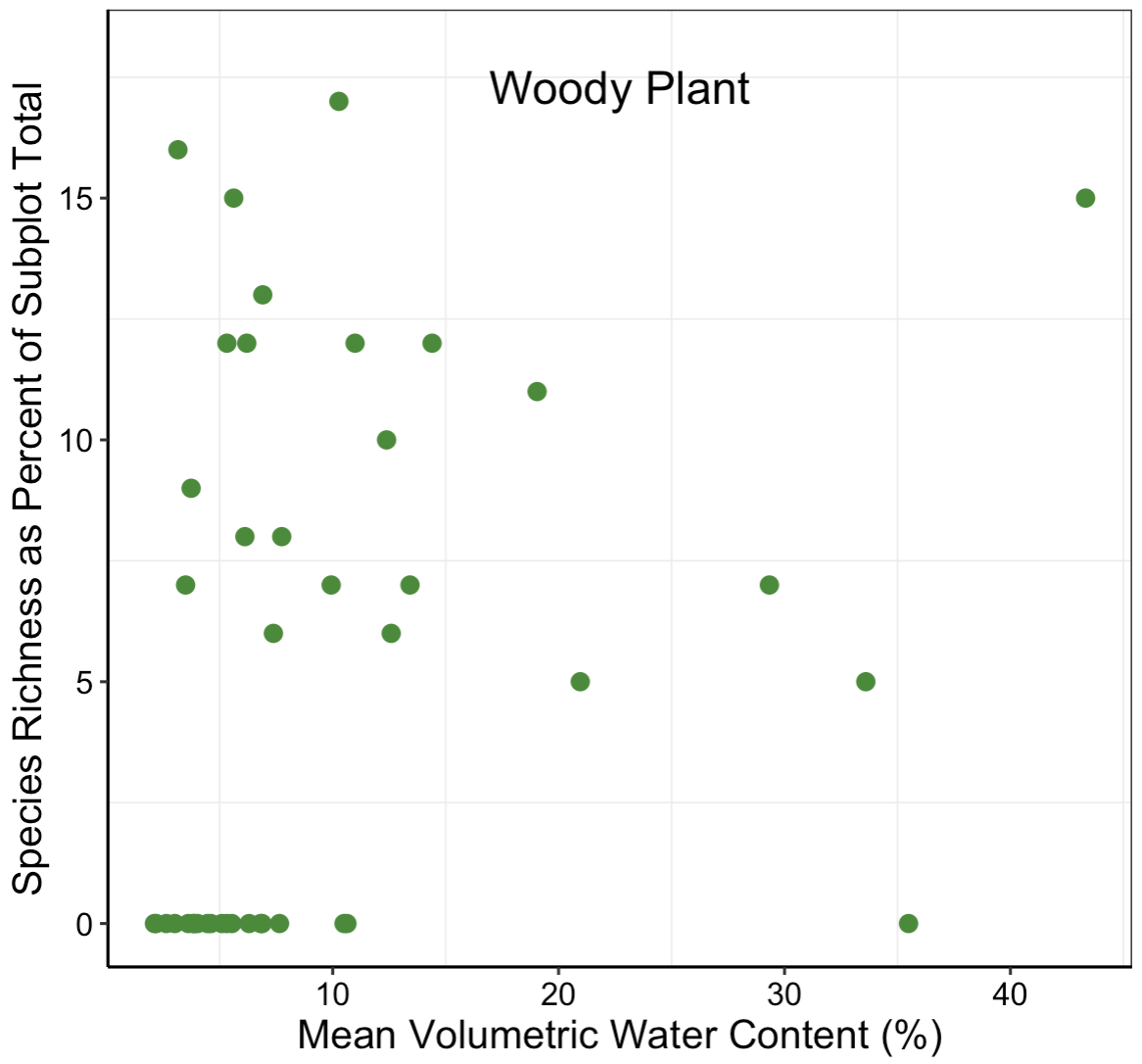

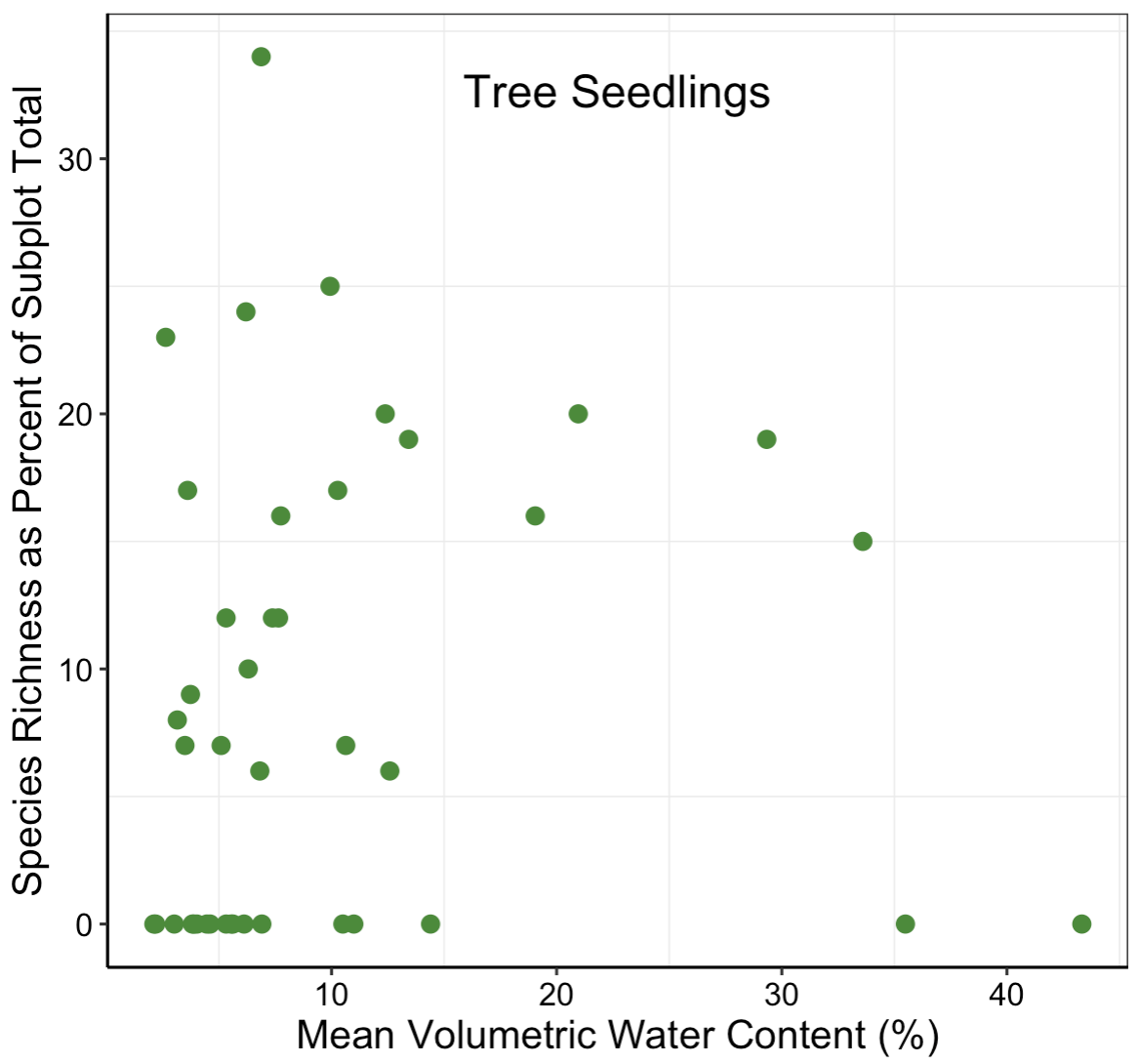

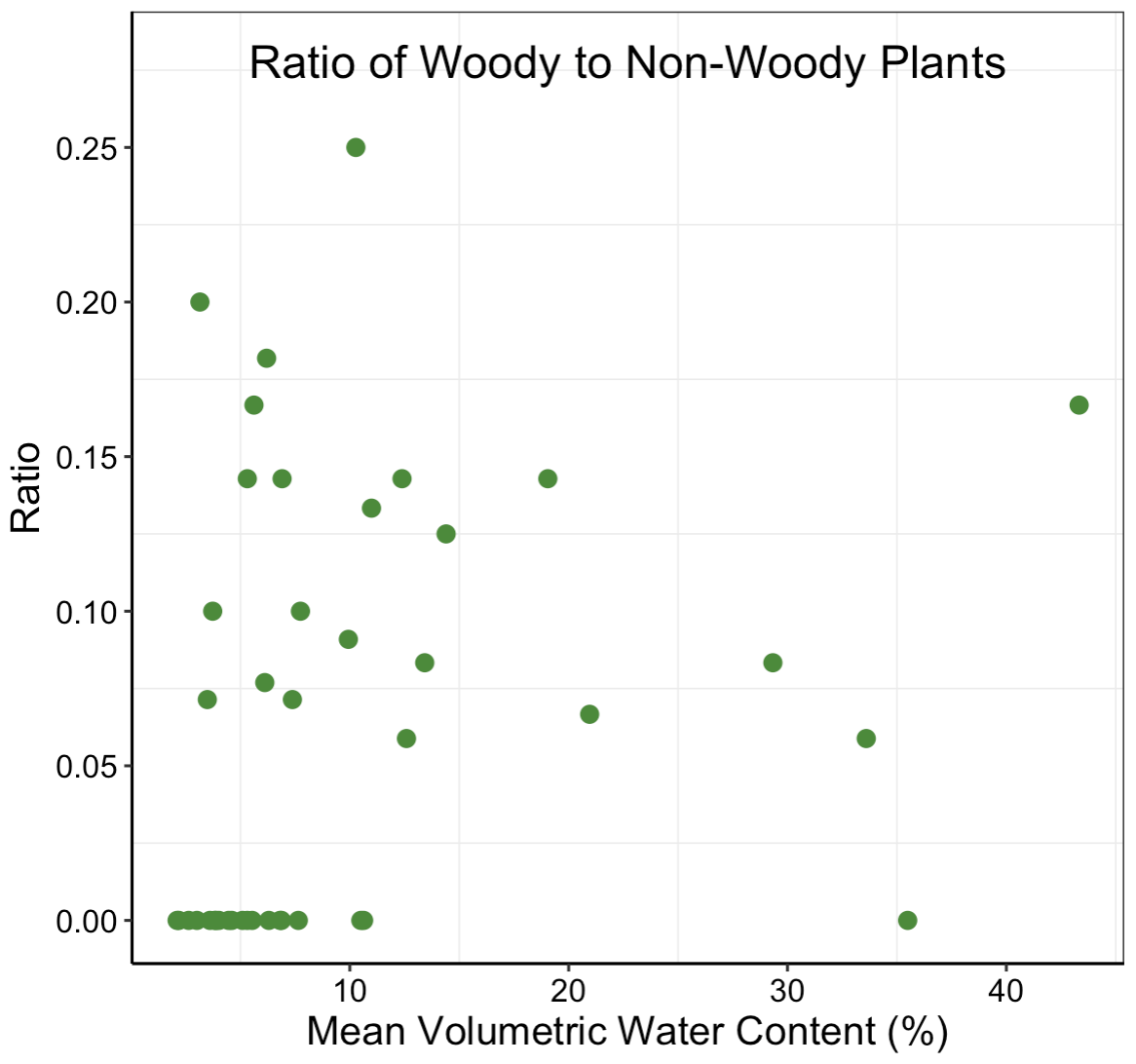

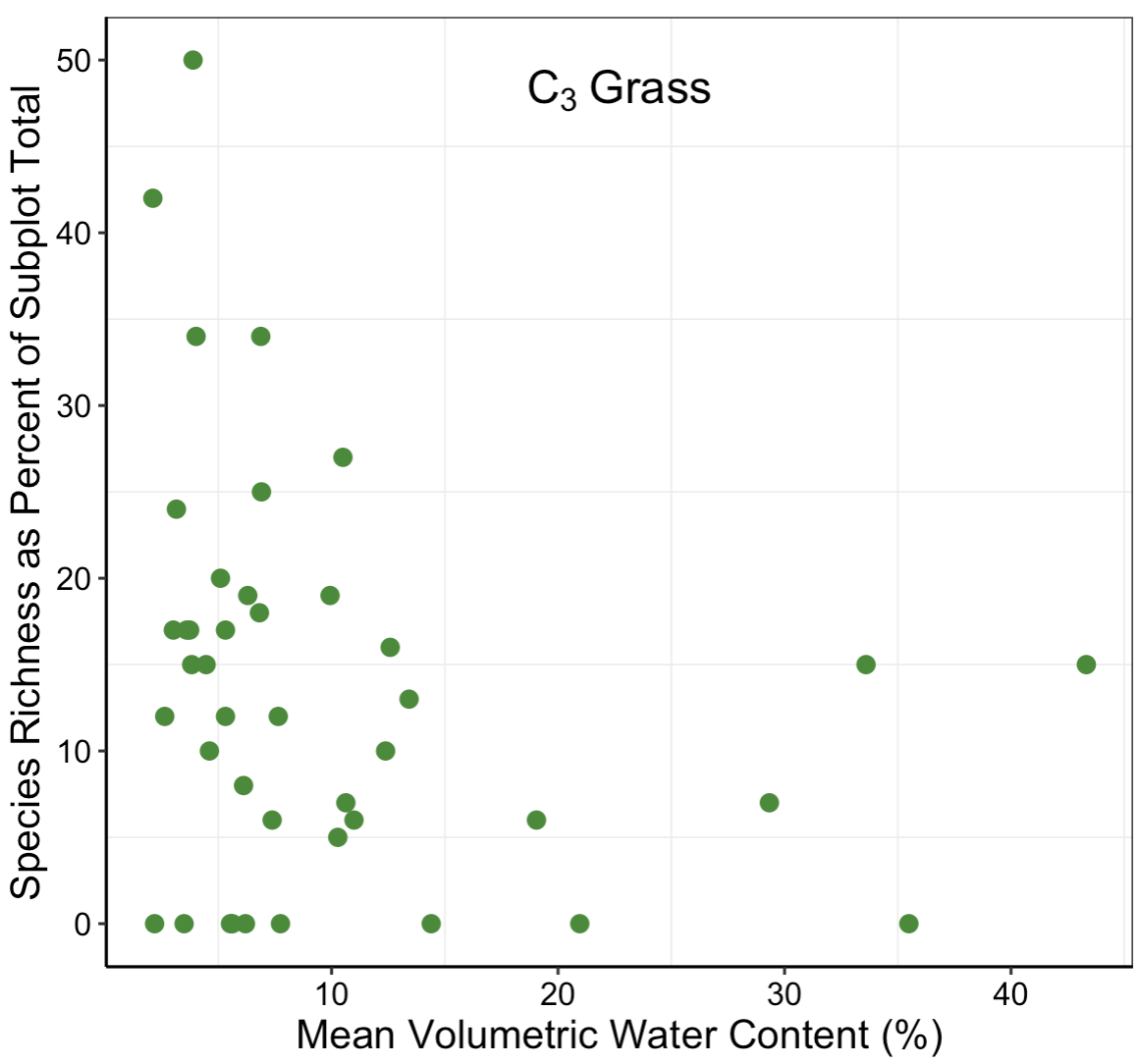

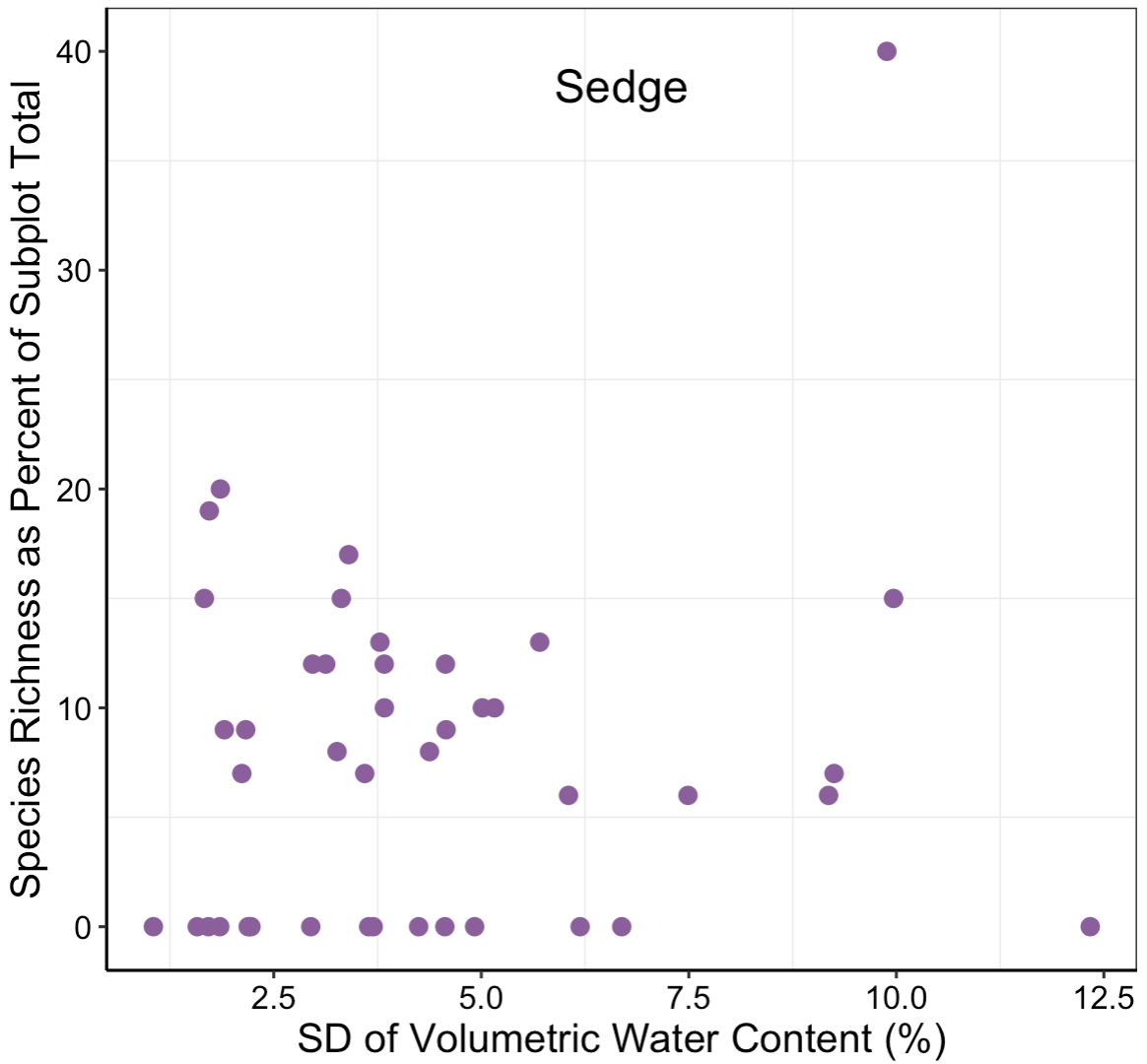

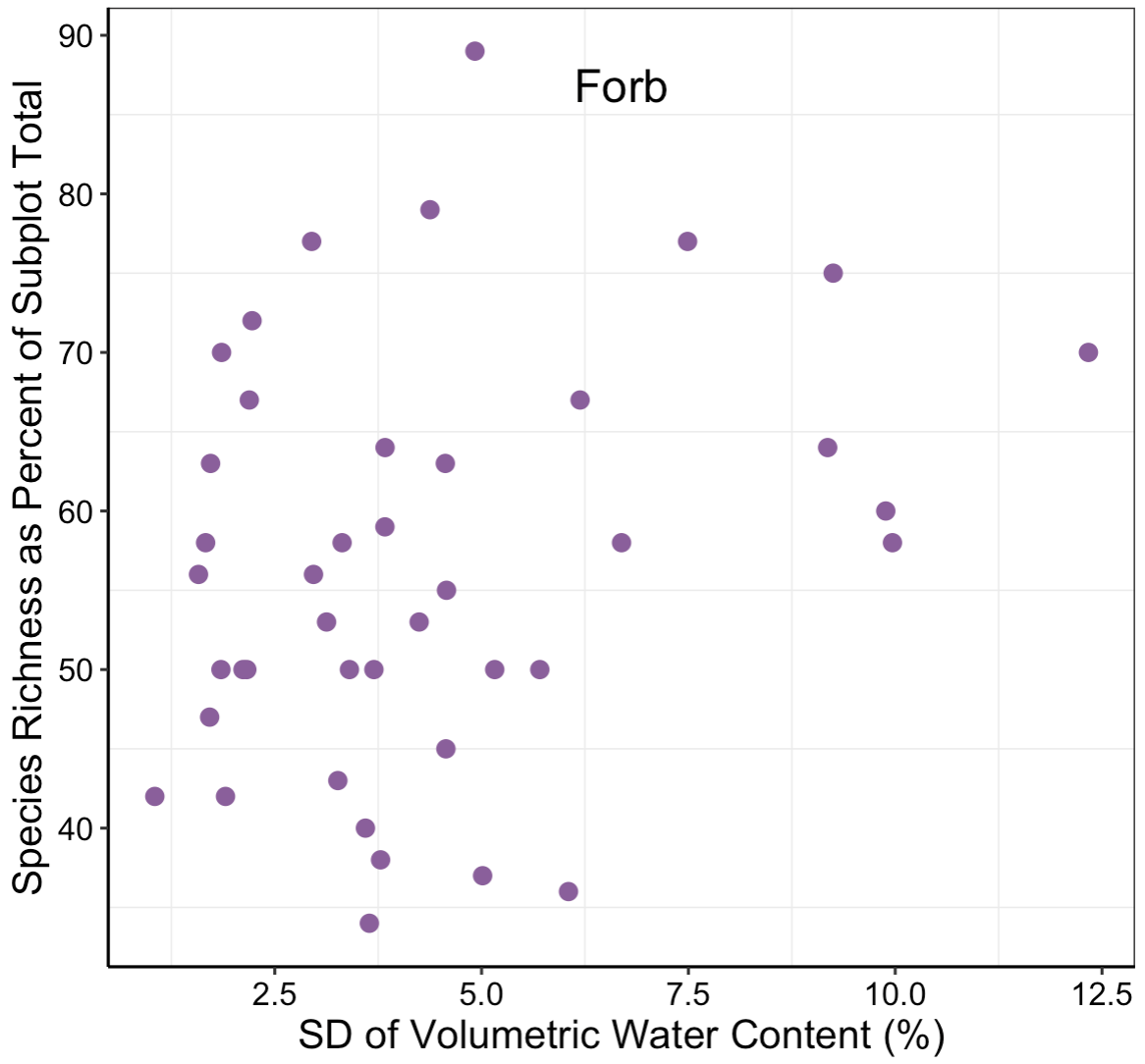

**(b)**

**(c)**

**(a)**

**(e)**

**(d)**

**(f)**

**(g)**

**Figure S5. Variation in understory woody and herbaceous species richness and the ratio of woody to non-woody species (rWNW) with respect to soil moisture in the Niobrara Plot in 2021.** Scatterplots show relationships from top-supported models (zero-inflated negative binomial models; binomial generalized linear model for forbs; Table 2) for species richness as a percent of the subplot total of (a) woody plants, (b) tree seedlings, (c) rWNW, and (d) C_3_ grasses versus mean volumetric water content (VWC); and of (e) C_4_ grasses, (f) sedges and (g) forbs versus variability (standard deviation, SD) in VWC. Point represents subplot values from the 42 herbaceous subplots for which VWC was measured.

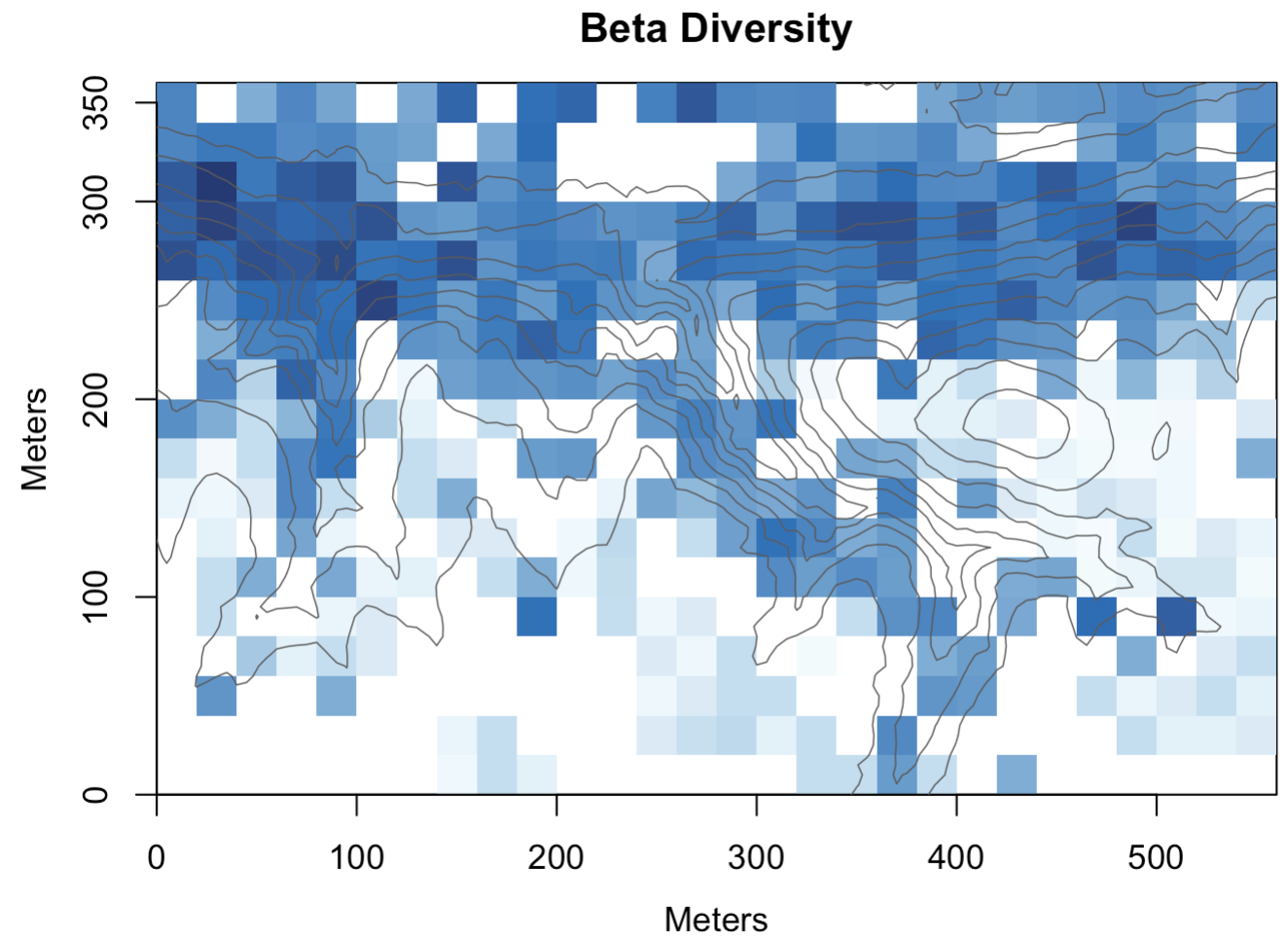

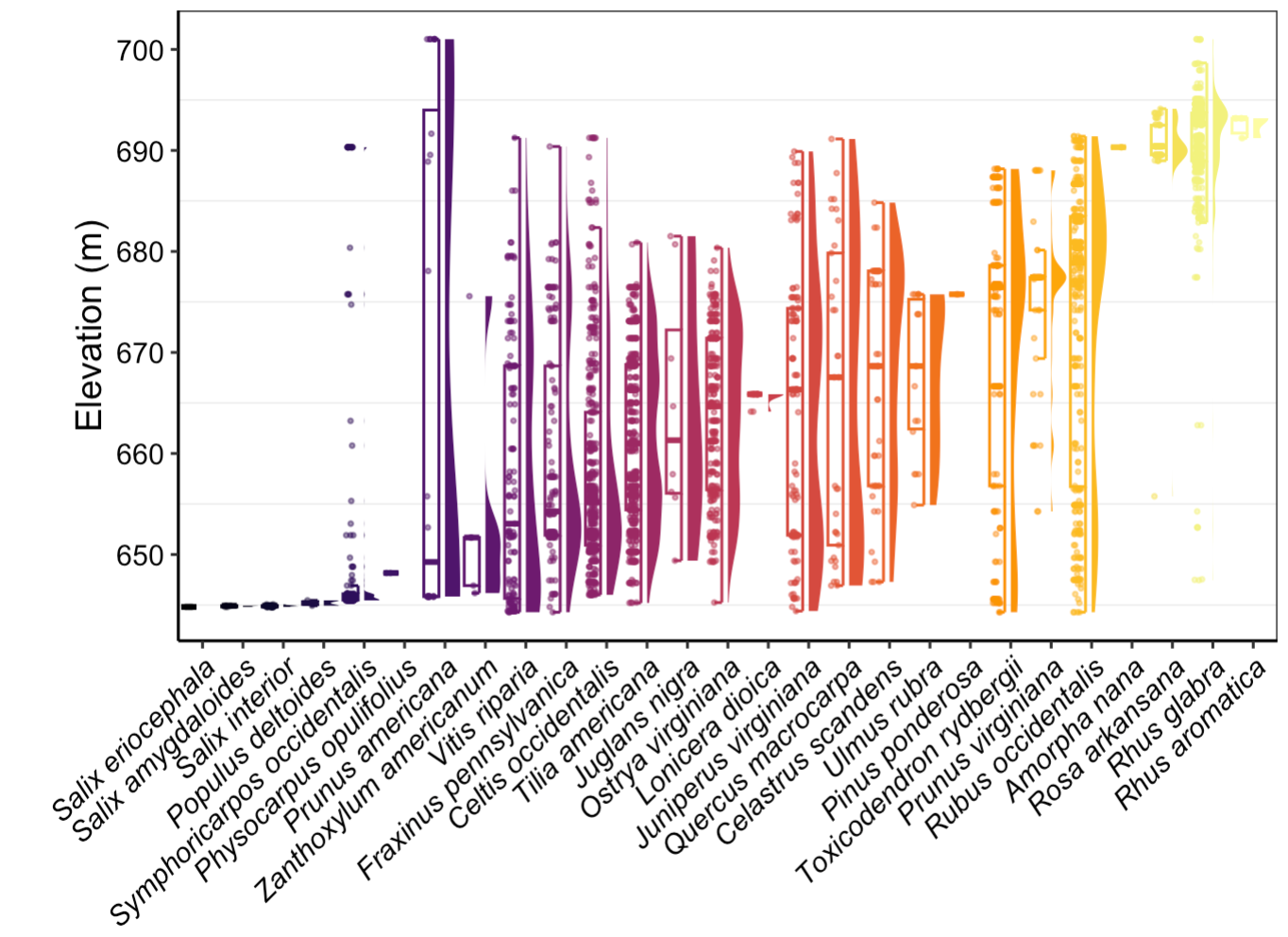

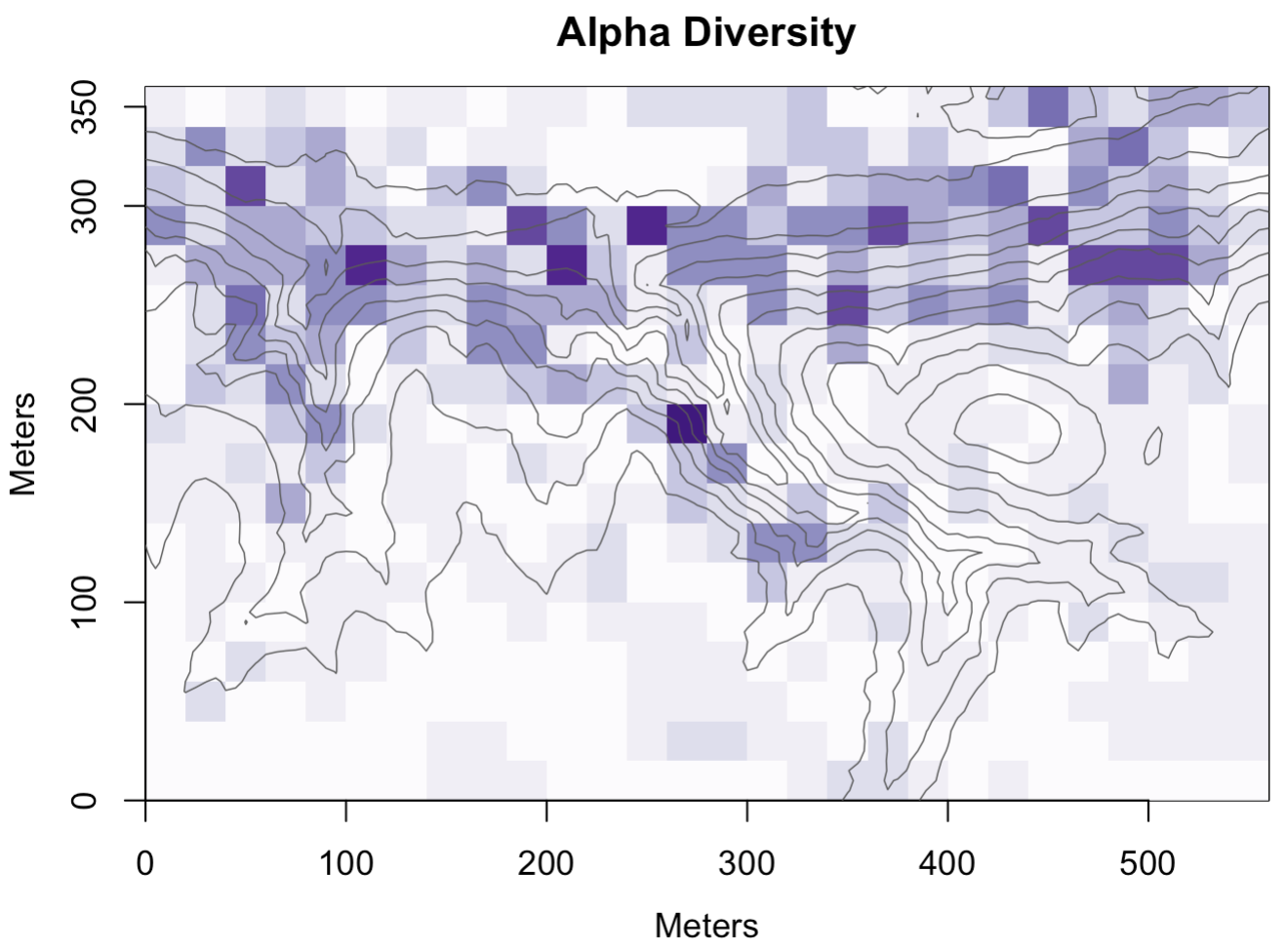

**(a)**

**(b)**

**(c)**

**Figure S6. Understory woody plant species turnover and alpha and beta diversity with respect to elevation in the Niobrara Plot in 2022.** (a) Combined box and violin plots showing the elevation distributions for woody species with >1 individual, ordered by increasing median elevation. Boxes indicate the median (center line) and first and third quartiles; lower and upper whiskers indicate the first and third quartiles ±1.5 times the interquartile range, and points indicate outliers; (b) heatmap showing species richness (alpha diversity) of 1 × 1 m subplots overlaid with 5-m elevation contour lines; 20 × 20 m quadrats are shaded from light to dark purple with increasing woody species richness (from 0-9 species) in the nested subplot; (c) heatmap showing variation in woody species composition (beta diversity) overlaid with 5-m elevation contour lines; quadrats are color-coded by scores from the first axis of a principal coordinates analysis of Cao dissimilarities in subplot-level species composition, with similarly shaded squares indicating more similar species assemblages. In (b) and (c), white quadrats represent nested subplots containing no woody stems <1 cm diameter at breast height.

**Table S7. Summary statistics for permutational analyses of variance testing the effects of habitat type and continuous topographic variation on understory woody plant species composition in the Niobrara Plot in 2021 and 2022.** Cao dissimilarities in species composition between subplot pairs were the response variable for each model. DF = degrees of freedom; significant effects at $\alpha$ = 0.05 shown in bold.

| **Habitat Type** | | | | | | |
| --- | --- | --- | --- | --- | --- | --- |
| **Year** | **Response Variable** | **Model Term** | ***F*-value** | **DF (Error, Total)** | ***R*^2^** | ***p*-value** |
| 2021 | Species composition | habitat type | 19.251 | 5, 355 | 0.216 | **0.001** |
| 2022 | Species composition | habitat type | 21.561 | 5, 346 | 0.240 | **0.001** |
| **Continuous Topographic Variation** | | | | | | |
| **Year** | **Response Variable** | **Model Term** | ***F*-value** | **DF (Error, Total)** | ***R*^2^** | ***p*-value** |
| 2021 | Species composition | slope | 25.452 | 1, 355 | 0.056 | **0.001** |
|  |  | elevation | 72.271 | 1, 355 | 0.160 | **0.001** |
|  |  | eastness | 0.336 | 1, 355 | 0.001 | 0.934 |
|  |  | northness | 2.621 | 1, 355 | 0.006 | **0.022** |
| 2022 | Species composition | slope | 29.731 | 1, 346 | 0.066 | **0.001** |
|  |  | elevation | 77.537 | 1, 346 | 0.171 | **0.001** |
|  |  | eastness | 0.608 | 1, 346 | 0.001 | 0.742 |
|  |  | northness | 3.402 | 1, 346 | 0.007 | **0.009** |

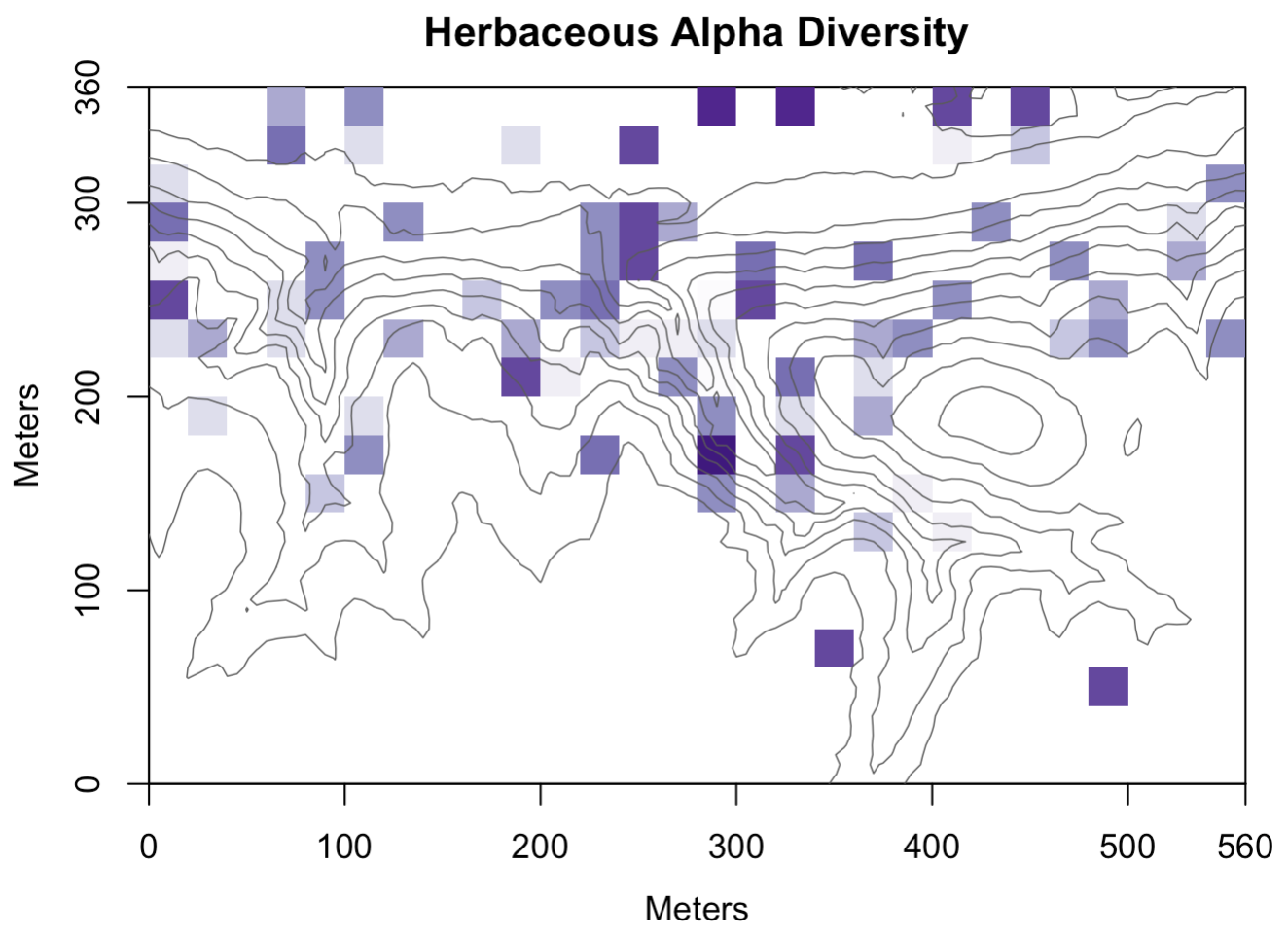

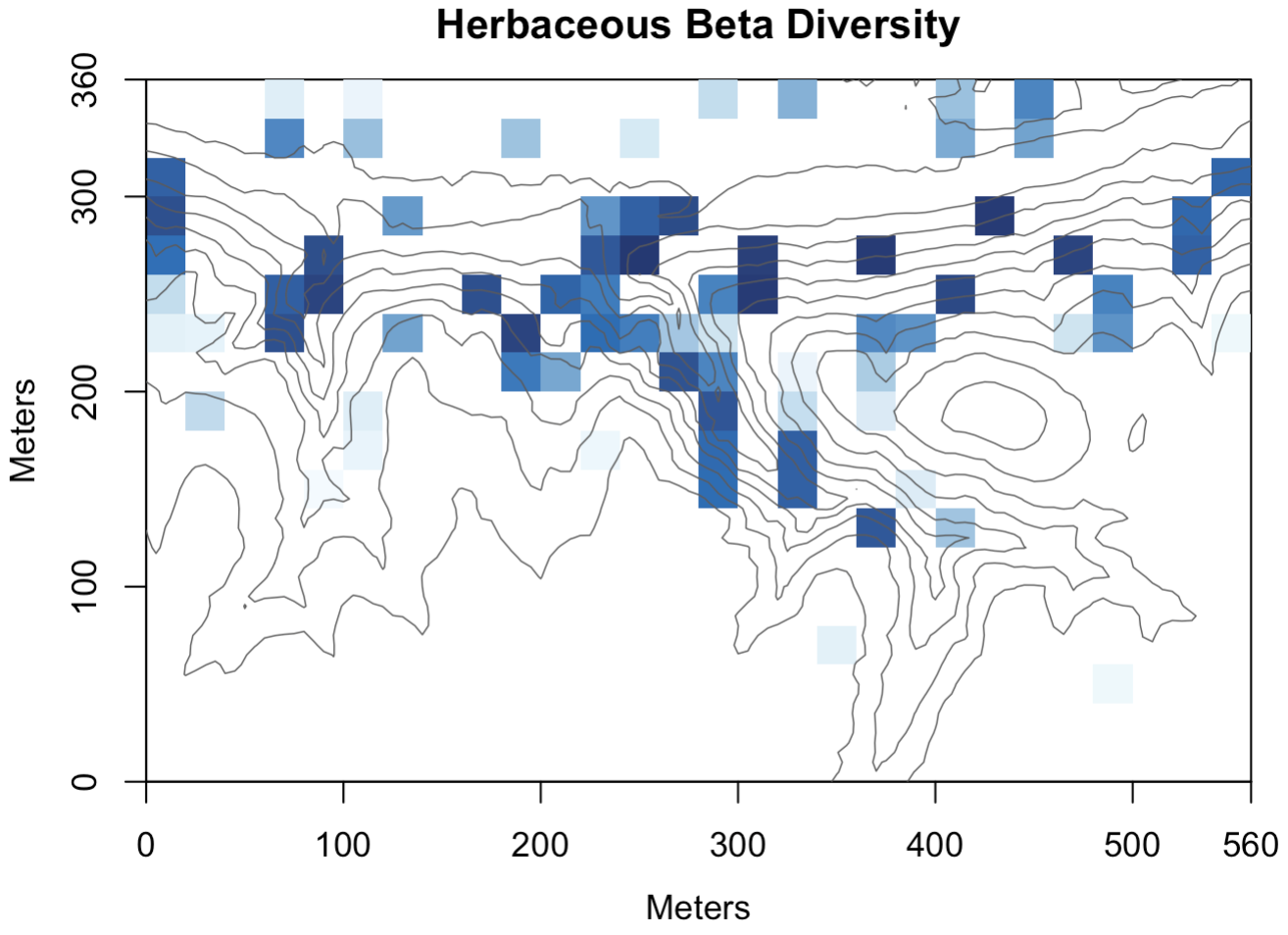

**(a)**

**(b)**

**Figure S7. Herbaceous plant alpha and beta diversity with respect to elevation in the Niobrara Plot in 2021.** (a) Heatmap showing alpha diversity of the 78 sampled 1 × 1 m subplots overlaid with 5-m elevation contour lines; 20 × 20 m quadrats are shaded from light to dark purple with increasing herbaceous species richness (from 2-19 species) in the nested subplot; (b) heatmap showing variation in herbaceous species composition (beta diversity) overlaid with 5-m elevation contour lines; quadrats are color-coded by scores from the first axis of a principal coordinates analysis of Sørensen dissimilarities (Bray-Curtis dissimilarity computed on presence/absence data) in subplot-level species composition, with similarly shaded quadrats indicating more similar species assemblages. In (a) and (b), white quadrats represent nested subplots that were not sampled for herbaceous plants.

**Table S8. Summary statistics for permutational analyses of variance testing the effects of habitat type and continuous topographic variation on herbaceous plant species composition in the Niobrara Plot in 2021.** Sørensen dissimilarities (Bray-Curtis dissimilarity on presence/absence data) in species composition between subplot pairs were the response variable for each model. Analyses were conducted across the 78 sampled herbaceous subplots spanning five habitat types (canyon bottoms, upland forest, upper canyon, floodplain, and ecotone). DF = degrees of freedom; significant effects at $\alpha$ = 0.05 shown in bold.

| **Response Variable** | **Model Term** | ***F*-value** | **DF (Error, Total)** | ***R*^2^** | ***p*-value** |
| --- | --- | --- | --- | --- | --- |
| Species composition | habitat type | 3.869 | 4, 75 | 0.179 | **0.001** |
| Species composition | slope | 7.871 | 1, 75 | 0.089 | **0.001** |
|  | elevation | 5.820 | 1, 75 | 0.066 | **0.001** |
|  | eastness | 1.351 | 1, 75 | 0.015 | 0.161 |
|  | northness | 2.172 | 1, 75 | 0.025 | **0.022** |
